# Detailed Social Network Interactions and Gut Microbiome Strain-Sharing Within Isolated Honduras Villages

**DOI:** 10.1101/2023.04.06.535875

**Authors:** Jackson Pullman, Francesco Beghini, Marcus Alexander, Shivkumar Vishnempet Shridhar, Drew Prinster, Ilana L. Brito, Nicholas A. Christakis

## Abstract

When humans assemble into face-to-face social networks, they create an extended environment that permits exposure to the microbiome of other members of a population. Social network interactions may thereby also shape the composition and diversity of the microbiome at individual and population levels. Here, we use comprehensive social network and detailed microbiome sequencing data in 1,098 adults across 9 isolated villages in Honduras to investigate the relationship between social network structure and microbiome composition. Using both species-level and strain-level data, we show that microbial sharing occurs between many relationship types, notably including non-familial and non-household connections. Using strain-sharing data alone, we can confidently predict a wide variety of relationship types (AUC ~0.73). This strain-level sharing extends to second-degree social connections in a network, suggesting the importance of the extended network with respect to microbiome composition. We also observe that socially central individuals are more microbially similar to the overall village than those on the social periphery. Finally, we observe that clusters of microbiome species and strains occur within clusters of people in the village social networks, providing the social niches in which microbiome biology and phenotypic impact are manifested.

---

The human microbiome plays a role in many aspects of human physical and mental health^1^, and the microbiome is in turn shaped by diverse factors. Diet, medications, lifestyle, and environmental exposures (such as animals) are known to affect a person’s microbiome composition^2–5^. But since few bacterial components of the human microbiome can survive for very long outside the human body, most such bacteria must somehow be acquired from other humans through diverse forms of physical contact. While maternal transmission to offspring is one obvious pathway^6–10^, adults may acquire normal flora from others beyond their mothers via social interactions^11^. Indeed, in models involving both wild mice and primates, gut microbiome information can predict a host’s social interactions, group membership, and network centrality^12–14^. And in human populations, recent evidence indicates the salience of household and spousal transmission of the microbiome^11,15^.

However, the influence of face-to-face social interactions beyond household contacts or closely related kin on the composition of the human microbiome is still incompletely understood^11,16^. Yet the impact of the broader set of social relationships that people have on their microbiome composition is surely relevant. Furthermore, the precise type of relationship (e.g., whether a person is a close friend, casual acquaintance, or neighbor), what the social interactions involve (e.g., in terms of face-to-face contact, sharing of meals, or physical greetings), and the frequency of interactions are likely also relevant. Here, we explore the impact on the microbiome, at the species and strain levels, of such detailed social network interactions within isolated rural villages, including interactions outside of households and with non-kin.

## Study Cohort

Microbiome acquisition patterns may be especially important to investigate in a traditional social setting involving face-to-face interactions within a circumscribed population that partakes of a traditional diet and is relatively devoid of antibiotics and other medications. Our cohort consists of 1,098 individuals in 9 isolated villages who are part of a larger population-based cohort we started following in 2015 for a different purpose^17^. The average distance from each of the 9 villages to the nearest other village among the 9 is 800 meters, and the average distance to the farthest other village is 12.3km. We combine face-to-face social networks mapped in detail for whole villages (i.e., sociocentrically); a comprehensive set of both individual and community-level characteristics regarding behavioral, socioeconomic, and health phenotypes; and detailed gut microbiome sequencing data. Such social network data from traditional populations are scarce^18–20^. Microbiome data from such developing world settings are also scarce, especially at large sample sizes^21^. The coverage rates (i.e., the percentage of people in the village-level social networks for whom microbiome samples could be collected) in these villages ranged from 43% to 76%. Village size ranged from 89 to 432 individuals; and the average household size was 4.46. The average age of participants was 40.9 (SD=17; range: 15-93); 62% were women; and 23.4% were married.

Several “name generator” questions were used to map the social networks, including questions such as “With whom do you spend free time?” and “Who do you trust to talk about something personal or private?” (See Table S1 for a summary of the name generator questions used). The total number of relationships identified in our cohort are as follows: Partner/Spouse (254), Father (209), Mother (369), Sibling (740), Child (262), Close Friends (915), Spend Free Time (1,061), and Personal or Private Conversation (1,129). Some of these relationships overlap, and, after network symmetrization, we identified 2,919 unique social network links. For individuals who report spending free time together, we also collected many details about the nature of these interactions, such as how often the pair spends free time together, whether they share meals, and how they typically greet each other.

## Microbiome Profiling

Strains can be very divergent, genetically and functionally, within a given microbe species^22,23^. Such variation can be very useful to bolster confidence that two people who have a similar strain acquired it from a common source. For instance, after filtering out strains associated with fermented foods, strain-sharing between two people can offer evidence that the shared strain resulted from direct interpersonal transmission rather than shared exposure to an environmental factor such as diet^24^. That is, documenting the same strain in two people can provide suggestive evidence for actual interpersonal transmission^23,25–28^.

We performed strain-level profiling with StrainPhlan 4 and detected putative transmission events between pairs of individuals. We summarize the strain-level similarity between two individuals with the strain-sharing rate metric which is equal to the number of shared strains between two samples divided by the number of species with available strain profiles that are present in both samples^29^. Overall, our data included information on 2,285 species and 183,195 strains (from the 682 species profiled by StrainPhlAn). We summarize the species-level beta diversity using one minus the Bray-Curtis dissimilarity of species’ relative abundances, and also with the Jaccard index when treating the presence or absence of a species as binary.

## Village-Level Microbiomes

Many villages in our setting tend to have distinctive microbiome patterns, some more than others (**Fig. 1**). Dimensionality reduction (with t-distributed stochastic neighbor embedding, t-SNE) of the centered log-ratio transformed species-level relative abundance data reveals clear differences in composition for most two-village comparisons and (to some extent) across the five most populous sites combined.

**Fig. 1.**
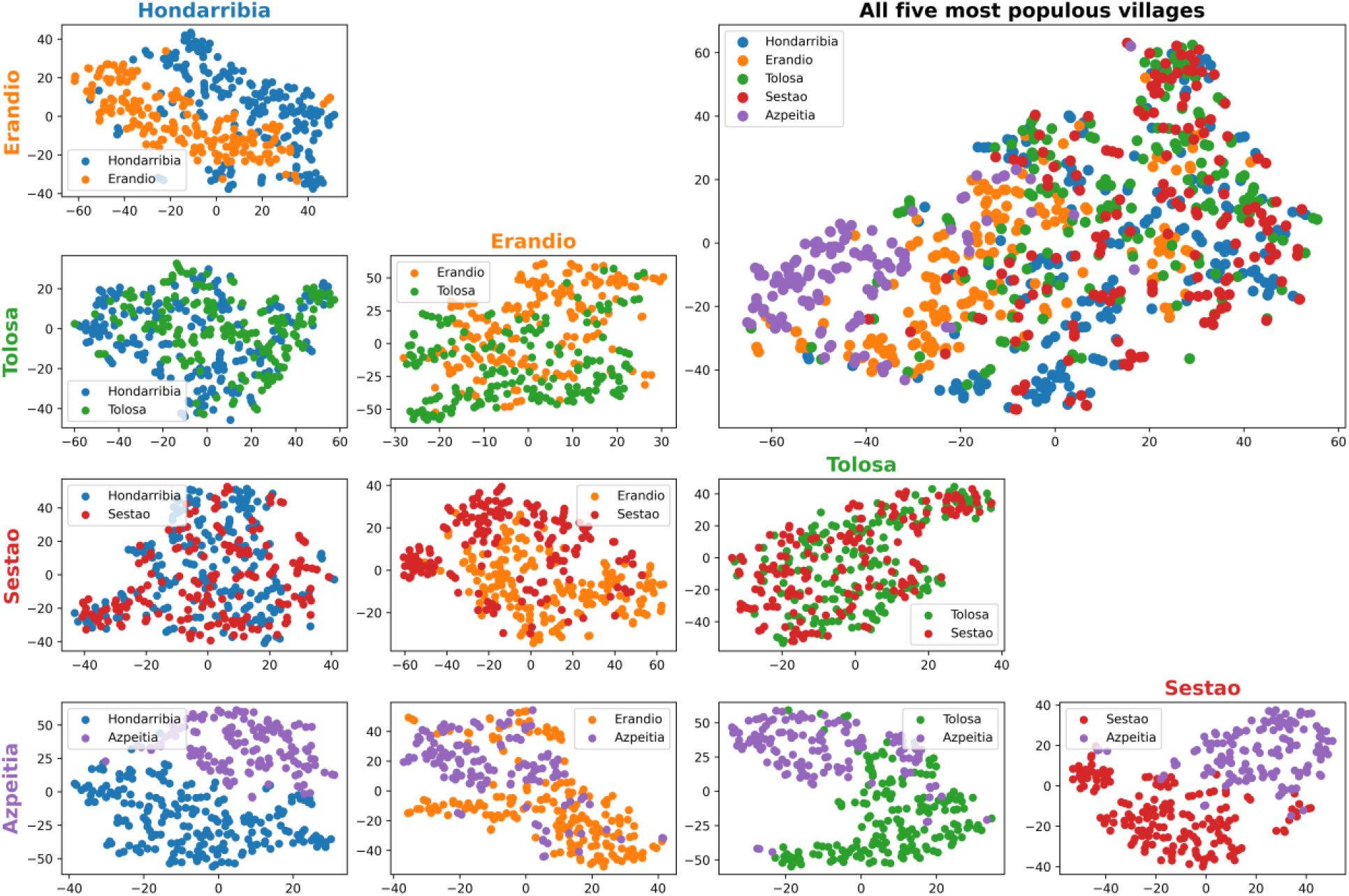
Visualization of microbiome species relative abundance data across villages. Data are shown after centered log-ratio transformation using t-distributed stochastic neighbor embedding (t-SNE), colored by village membership, for the five most populous villages in the Honduras microbiome cohort. A two-dimensional t-SNE plot for all five of the most populous villages combined is in the top right, and the remaining plots show the two-dimensional t-SNE visualizations for all distinct pairs of the five most populous villages. Microbiome samples are clearly distinguished by village membership for most pairs of villages and to some extent still when all five villages are combined. The distinction of microbiome clusters by village appears to depend on the village; for example, the village Azpeitia maintains a relatively distinct cluster in all visualizations.

## Strain Sharing Across Multiple Relationship Types

We observe microbiome species-sharing and strain-sharing occurring across many relationship types. Pairs of individuals with diverse sorts of relationships (partner/spouse, father, mother, sibling, child, close friend, free time, personal-private) share significantly more strains with each other than other pairs of people from within the same village with no relationship, and we observe a social-distance-based gradient of strain-sharing among relationships (two-sided Wilcoxon rank-sum tests, max *P_adj_* ≤ .05) (**Fig. 2A**). Spouses and relationships between individuals living in the same building have the highest levels of strain-sharing (median strain-sharing rate of 13.5% and 12.9% respectively). While prior studies have revealed potential household and familial transmission^11,15,24^, we also observe a significantly elevated strain-sharing rate between non-kin relationships living in different houses (median 6.2%, permutation *P* < 2.2 × 10^-16^). We observe minimal amounts of strain-sharing between individuals living in the same village *without* a social relationship (median 3.9%), potentially a result of shared village environments or network-wide circulation of strains. And we observe a low strain-sharing rate between individuals living in different villages (median 2%).

**Fig. 2.**
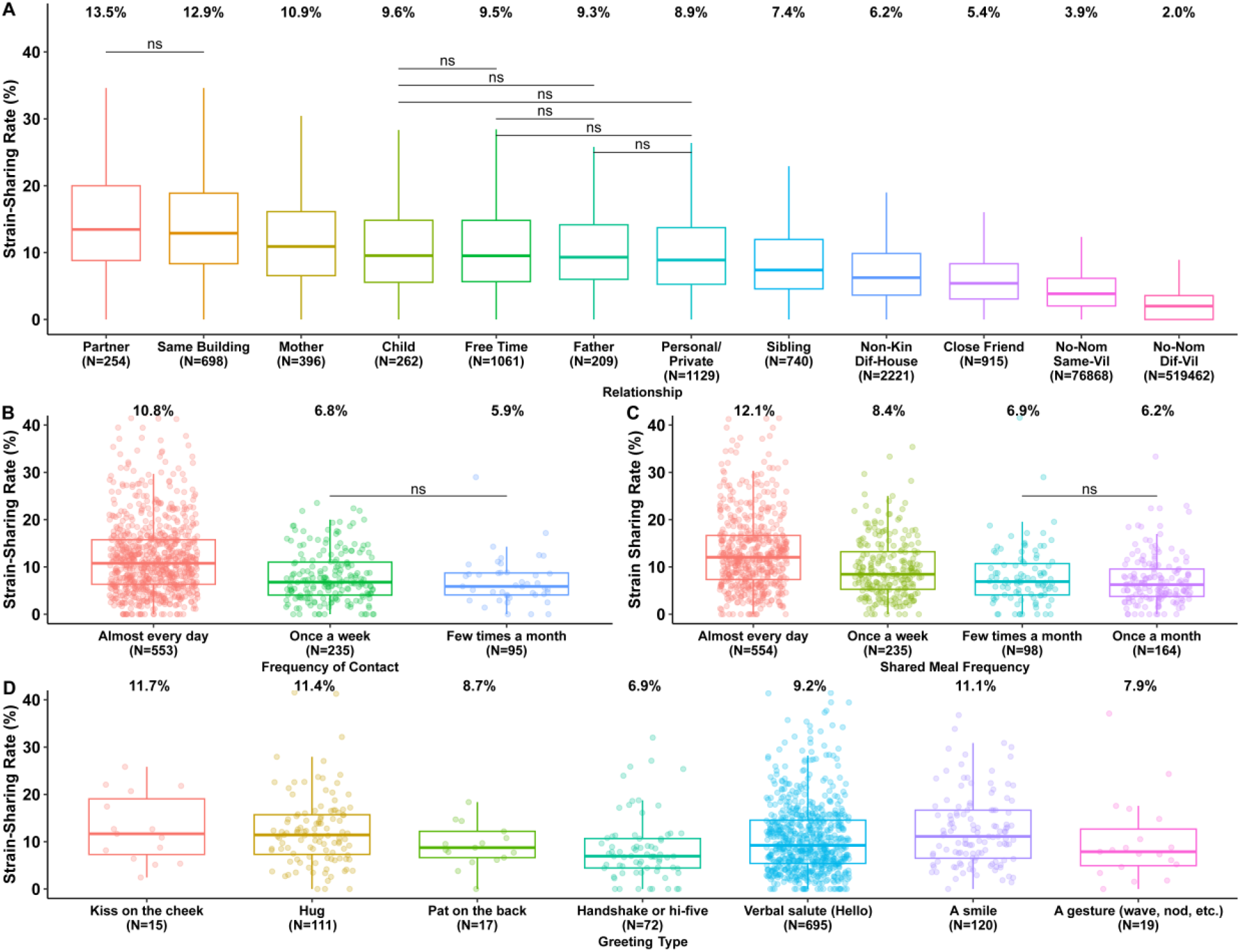
Strain-Sharing Across Multiple Relationship Types. **(A)** Distribution of strain-sharing rates based on relationship type. All pairwise relationship comparisons are significantly different, except for those marked ‘ns’ (Kruskal–Wallis test, χ^2^ = 43,416, n = 604,215, *P* < 2.2 × 10^-16^, two-sided Wilcoxon rank-sum tests, NS, not significant (*P_adj_* ≥ .05)). The final two boxes quantify the strain-sharing rates between all pairs of individuals living in the same village without a nominated relationship, and all pairs of individuals living in different villages, respectively. Median values for each distribution are printed at the top of each box. **(B)** The propensity to share strains increases as a function of how often a pair spends free time together. Only the non-significant pairwise comparisons are indicated (Kruskal–Wallis test, χ^2^ = 62.3, n = 1,033, *P* = 3.0 × 10^-14^, two-sided Wilcoxon rank-sum tests; NS, not significant (*P_adj_* ≥ 0.05)). **(C)** The propensity to share strains increases as a function of how often a pair shares meals together. Only the non-significant pairwise comparisons are shown (Kruskal–Wallis test, χ^2^ = 122.4, n = 1,051, *P* < 2.2 × 10^-16^, two-sided Wilcoxon rank-sum tests; NS, not significant (*P_adj_* ≥ .05)). **(D):** The strain-sharing rate varies by the typical way individuals greet each other (Kruskal–Wallis test, χ^2^ = 23.3, n = 1,049, *P* = 0.0007).

Since species distributions are to some extent village-dependent (**Fig. 1**) and individuals in the same village have higher strain-sharing rate than individuals living in different villages (**Fig. 2A**), village-level sharing can serve as a baseline for comparison. To account for both the potential influence of village-level microbiome niches and of village-level network structure, we compared each relationship distribution to 100 samples from a within-village relationship permutation (e.g., swapping mother-child pairs in the same village; see Methods), and the same pattern of variation in strain-sharing by relationship type is observed (Fig. S1). This result is also observed at the species level (Fig. S2, S3), although to a lesser extent, possibly suggesting that strain-sharing is more likely to be a result of direct transmission than species-level sharing, which could potentially originate from multiple sources (such as a shared environment).

For individuals who report spending free time with someone else, we also examined how microbiome strain-sharing may relate to how often they spend free time together, how often they share meals together, and how they typically greet each other (**Fig. 2, B, C, and D**). The frequency that a person spends free time with someone, whether casually or through a meal, is associated with an increase in their propensity to share strains (Free-time, Kruskal–Wallis test, χ^2^ = 62.3, n = 1,033, *P* = 3.0 × 10^-14^; Meals, Kruskal–Wallis test, χ^2^ = 122.4, n = 1,051, *P* < 2.2 × 10^-16^). This result holds even when excluding the effect of living in the same house and kinship (Free-time, Kruskal–Wallis test, χ^2^ = 12.7, n = 574, *P* = .0017; Meals, Kruskal–Wallis test, χ^2^ = 22.4, n = 586, *P* = 5.4 × 10^-5^) (Fig. S4), suggesting that close physical proximity and shared meals are potential transmission routes when individuals are not cohabiting. Diet is a key modifiable factor influencing the composition of the human microbiome^30^ and shared foodstuffs are a potential explanation for microbial sharing. That is, shared meals can lead to similar gut microbiomes among group members because eating similar foods at the same time can lead to microbial sorting in the gut, creating similar microbial communities even if there is no direct exchange of microbes between individuals^31^.

The strain-sharing rate for non-kin living in different houses who spend free time together almost every day (median of 7.8%) is higher than the strain-sharing rate for friends who see each other only once a week (6.5%) or a few times a month (5.7%, Fig. S4). A similar gradient is observed with the frequency that non-cohabiting non-kin share meals together, with those sharing meals daily or weekly (median strain-sharing rates 8.5% and 8.0% respectively) sharing more than pairs who share a meal together a few times or only once a month (6.7% and 5.8%). Physical contact via kissing, hugging, and touching can also result in the transfer of microbes^32^ and studies have shown that the oral microbiome is influenced by different greeting types, such as kissing^33^. Pairs who greet each other with a kiss on the cheek have the highest median strain-sharing rate (median 11.7%) – although, perhaps due to the low sample size and diversity of greeting types mentioned (see Methods), the strain-sharing rates across most greeting types are not significantly different from each other (**Fig. 2D)**. We had hypothesized that individuals reporting a physical greeting type would have a higher strain-sharing rate than those who self-report using a non-physical greeting type, but we did not observe this (Fig. S5).

We find that mothers tend to have a significantly higher strain-sharing rate with their children than fathers (two-sided Wilcoxon rank-sum test, *P_adj_* ≤ 0.05) (Fig. S6). Previous research has shown that mothers may transmit bacterial strains to children during childbirth^6^, and, although our study does not contain infants (the youngest participant is 15), this higher strain-sharing rate may be a result of the retention of strains transmitted during infancy (we observe a higher strain-sharing between mothers and their children the younger the child is; see Fig. S6). Alternatively, the high mother-child strain-sharing rate may be explained by cultural factors that result in more opportunities for household transmission events between mothers and children relative to fathers and children, since current Honduras gender roles are strict and mothers tend to be at home with their children during adolescence more than fathers. At the species level, while mothers have a higher average Bray-Curtis similarity to their children than fathers, the trend is insignificant, suggesting some of the effects of mother-child microbiome transmission may have a stronger signal at the strain level than species level (Fig. S6). As children age, they may have a higher propensity to populate their microbiome with novel species from non-maternal sources, which could lower species-level mother-child similarity yet still maintain a high strain-sharing rate due to maternal strains retained from infancy.

In contrast to past analyses^11^, we find no evidence that women are more likely to share strains with their direct social connections than men (two-sided Wilcoxon rank-sum test, *P_adj_* ≥ 0.05) (Fig. S7). In fact, at the species level, we observe the opposite trend, where men are more microbially similar to their direct connections than women are, based on Bray-Curtis similarity (two-sided Wilcoxon rank-sum test, *P_adj_* ≤ 0.05, Fig. S7). A large portion of this effect appears to stem from brothers having more similar microbiomes to each other than sisters (median Bray-Curtis similarity 0.39 and 0.284 respectively; two-sided Wilcoxon rank-sum test, *P_adj_* ≤ 0.05; see Fig. S7). This same effect does not appear with the Jaccard index, suggesting the absolute difference in species between brothers and sisters is not large, but that sisters are more variable in their relative abundances than brothers. The contrast with prior work may relate to different social habits in Honduras compared with other places where this has been studied (such as Fiji^11,16^) or with differences between the oral and gut microbiome.

## Strain-Sharing Strongly Predicts Social Relationships

To evaluate the strength of strain-sharing and species-sharing across relationship types, we implemented a mixed-effects logistic regression model to predict whether any pair of individuals in a village has a social or familial relationship. If there is a strong imprint of the social network in the microbiome network, we would expect the microbiome similarity between two individuals to be a strong predictor of a social relationship. We created a second model to test the predictive power of microbial-sharing outside of the family and household by removing kin and household relationships from our positive class. To account for potential confounding by socio-demographic factors, we created three versions of each predictive model, one with strain-sharing rate as the only predictor (in addition to a random slope for each village); one including age and gender; and one including the socio-demographic variables age, gender, wealth, education, religion, and indigenous status (See Methods).

Using strain-sharing rate as the only predictor, the classifier achieves moderately strong performance across all relationships and also in non-kin different-household relationships (AUC 0.73 ± 0.007 and AUC 0.67 ± 0.009 respectively, **Fig. 3 A** and **3B**. **Fig. 3C** and **3D** show respective model predictions for the village of Ermua). Species-level similarity, as measured by Jaccard and Bray-Curtis similarity, achieves poor performance (All Relationships; Jaccard, AUC 0.55 ± 0.008, Bray-Curtis, AUC 0.54 ± 0.008 – Fig. S8).

**Fig. 3.**
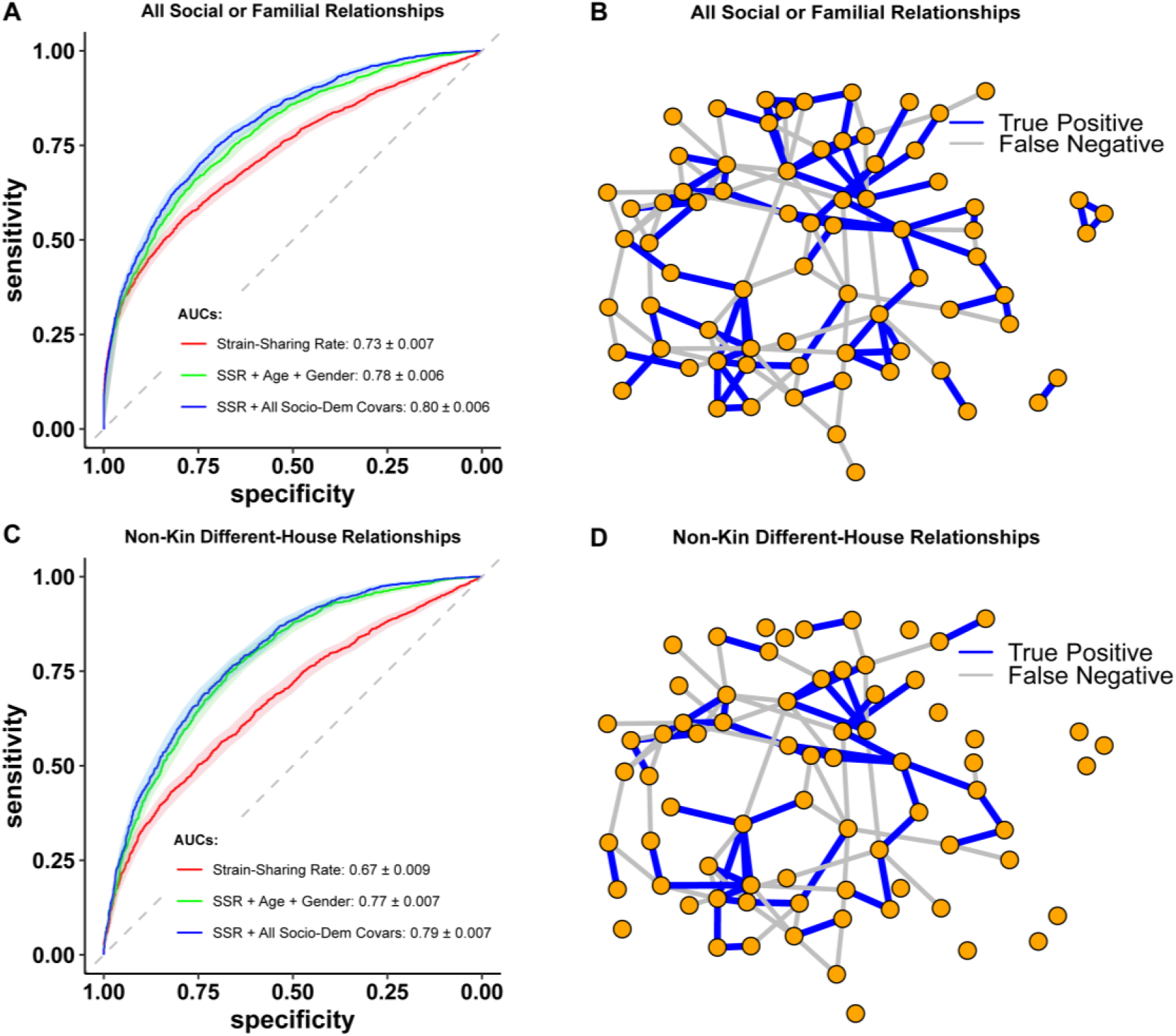
Strain-level Model Predicting Social Connections. Area under ROC curves predicting a social or familial relationships (**A)** or only non-kin, different-house relationships (**B**) compared to un-nominated pairs living in the same village; 95% DeLong confidence intervals are shaded surrounding each line. The diagonal dotted line indicates an AUC of 0.50, a “test” no better than chance. Legends report the means and standard deviations for each classifier’s AUC. SSR: Strain-Sharing Rate. **(C and D)** True-positive and false-negative network predictions for all relationships and non-kin different-house relationships for the village of Ermua. The model including all relationships performs better than the non-kin, different-house model as there is increased sharing within households and amongst kin. The model generally performs better at predicting within social clusters as there is increased strain-sharing around nodes with a high clustering coefficient.

To understand how much more strongly strain-sharing indicates a social relationship compared to socio-demographic attributes in our model, we use permutation feature importance metrics (see Methods). The permutation feature importance is defined to be the decrease in a model score when a single feature value is randomly shuffled^34^. This method shows that the strain-sharing rate is a stronger predictor of a relationship in both models than similarity along any other individual socio-demographic dimension in our study (Fig. S9).

## Geodesic Horizon of Microbiome Similarity

Observed strain-sharing patterns in villages could be consistent with possible chains of transmission. For instance, if someone’s microbiome is more similar to their friends than would be expected if the microbiome distribution and social network were independent, are they also more similar to their friends of friends? By measuring complete village social networks, we can evaluate how far into a network microbiome strain-sharing extends. We can calculate the distribution of the strain-sharing rate by the geodesic distance (shortest path between the vertices in a social network) between any two individuals. Under the null hypothesis that a host’s social network has no marginal relationship with their own microbiome composition, we can construct a permutation null distribution by randomly swapping the microbiome of every individual in a village and observing the null distribution of strain-sharing rate by geodesic distance. As previously noted, first-degree relationships have a much higher strain-sharing rate than we would expect under the null hypothesis (median 7.5%). But this effect also extends to second-degree connections (4.7%) before falling off at a social horizon of third-degree connections (4.1%), where pairs of people have a median strain-sharing rate no higher than would be expected under the null hypothesis within these small villages (**Fig. 4A**) (See Fig. S10 for species-level analyses).

**Fig. 4.**
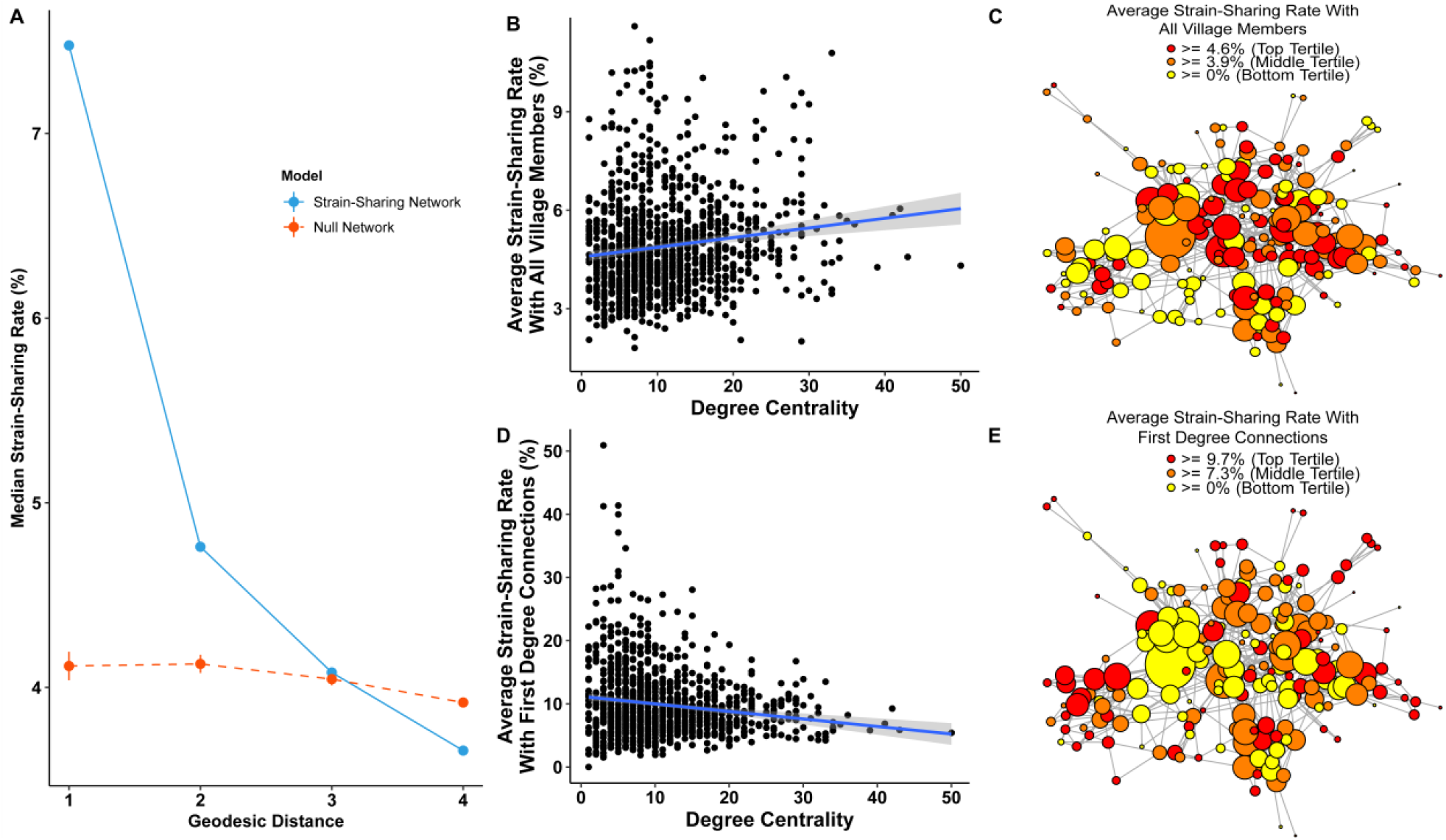
Strain Sharing and Social Network Position. **(A)** The strain-sharing rate by geodesic distance is shown. Null distributions were calculated based on 10,000 random samples from a within-village microbiome permutation. The null distribution slopes downwards because of the effect of large networks; larger villages have more pairs with a higher geodesic distance between them and, on average, have a lower average strain-sharing rate between individuals (see Fig. S11 for more details); 95% Confidence intervals are plotted around the null distributions. **(B)** As an individual’s degree centrality (number of social connections) increases and they get more socially embedded in the village, their average strain-sharing rate with the village also tends to increase. (**C)** Example social network for Hondarribia. Individuals with more connections (degree) and a higher network centrality (betweenness or eigenvector) tend to be more microbially typical as well. Nodes are colored according to their average strain-sharing rate with the rest of the village, and nodes are sized according to their degree centrality. (**D)** When individuals have a wider variety of social connections (increasing degree centrality), between-host heterogeneity tends to increase, and individuals on average have a lower strain-sharing rate with their first-degree connections. **(E)** Example social network for Hondarribia. Central individuals with a wider variety of social connections on average have a lower strain-sharing rate with their first-degree connections. Nodes are colored according to their average strain-sharing rate with their first-degree connections, and nodes are sized according to their degree centrality.

## Social Network Position and Strain Network Position

The strain-sharing patterns we have seen allow us to view the possibility of microbiome transmission from the framework of disease ecology. Individuals who are more socially central may also be more microbially central and more exposed to strains potentially spreading within a network. We may expect that socially central individuals are more microbially related to the rest of the village and more representative of the “social microbiome,” or the microbial metacommunity of transmittable strains within the village^14^. After controlling for sociodemographic covariates (see Methods), we tested whether there was a relationship between an individual’s average strain-sharing rate with all others in the village, representing their microbiome centrality, and their social network centrality. We examined three measures of social network centrality: degree centrality (the number of connections); normalized betweenness centrality (the fraction of the number of shortest paths in a network that pass through a person); and eigenvector centrality (a function of the number of connections and the centrality of those connections).

All measures of social network centrality were positively correlated with an individual’s average strain-sharing rate to the rest of the village, suggesting that the microbiome of more socially central people is more representative of the social microbiome in the village (linear mixed effects regression, n =1036; Degree, *β* = .017, *P* = 0.00044; Normalized Betweenness, *β* = 2.8, *P* = 0.0094; Eigenvector, *β* = .55, *P* = 0.0018) (**Fig. 4B**). This effect is apparent visually in **Fig. 4C** where individuals are colored based on their average strain-sharing rate with the rest of the village, with red individuals sharing more and yellow individuals sharing less. More socially central individuals tend to have higher strain-sharing rates with everyone else in the village than socially peripheral individuals.

We may also hypothesize that, while socially central individuals are *more* microbially representative of the overall network possibly as a result of the greater quantity of social contact, they may be *less* microbially similar to their typical first-degree social connection. An increase in the number and frequency of social contacts globally within the village may lead to higher between-host heterogeneity in gut microbial diversity locally. A very popular individual may be more representative of the social group at large, but, as a result of their many social interactions, they may be more removed from each of their individual connections, in a paradox of popularity. Indeed, we observe that increases in degree, eigenvector centrality, and betweenness centrality all correlate with a decrease in average microbiome similarity to first-degree connections. (linear mixed effects regression, n =1036; Degree, *β* = −.089, *P* = 0.00014; Normalized Betweenness, *β* = −19.5, *P* = 0.00019; Eigenvector, *β* = −2.6, *P* = 0.0014) **(Fig. 4D**). Gregarious individuals are less intimately related to their social connections, whereas individuals with only a handful of social connections tend to be more intimately microbially tied to those connections. This effect is apparent visually in **Fig. 4E** where individuals are colored based on their average strain-sharing rate with their first degree connections, with red individuals sharing more and yellow individuals sharing less. More socially central individuals tend to have lower average strain-sharing rates with their first degree connections, whereas peripheral individuals share much more (this trend is the opposite of **Fig. 4C**, with the red and yellow colorings generally being reversed).

## Social Clusters and Microbiome Clusters

The strain-sharing patterns we have seen along village, household, familial, and social lines would mean that social clusters (i.e., communities of more densely interconnected individuals within the village networks) should also have shared sets of particular microbiome strains^14^. That is, the phenomena so far documented should come to instantiate or to reflect niches of microbiomes within niches of people (analogous in some ways to soil biology^35^).

At the smallest scale, individuals with a higher clustering coefficient (i.e., transitivity – that is, the fact that three individuals are connected in a triangle) are more likely to have a higher average strain-sharing rate to those connections (linear mixed effects regression, n =1036, *β* = 2.97, *P* = 6.46 x 10^-6^). Having relationships to others who are also connected may facilitate microbiome circulation, instantiating microbiome niches within social niches. Individuals with a high clustering coefficient (≥.75) who form small-world networks with their connections have a high average strain-sharing rate with their first-degree connections (median 9.68%). Conversely, individuals with a low clustering coefficient ( ≤ .25) (and thus links to disparate parts of the network) have a lower average strain-sharing rate with their first-degree connections (8.05%) (two-sided Wilcoxon rank-sum test, *P_adj_* ≤ .05) (**Fig. 5A)**.

**Fig. 5.**
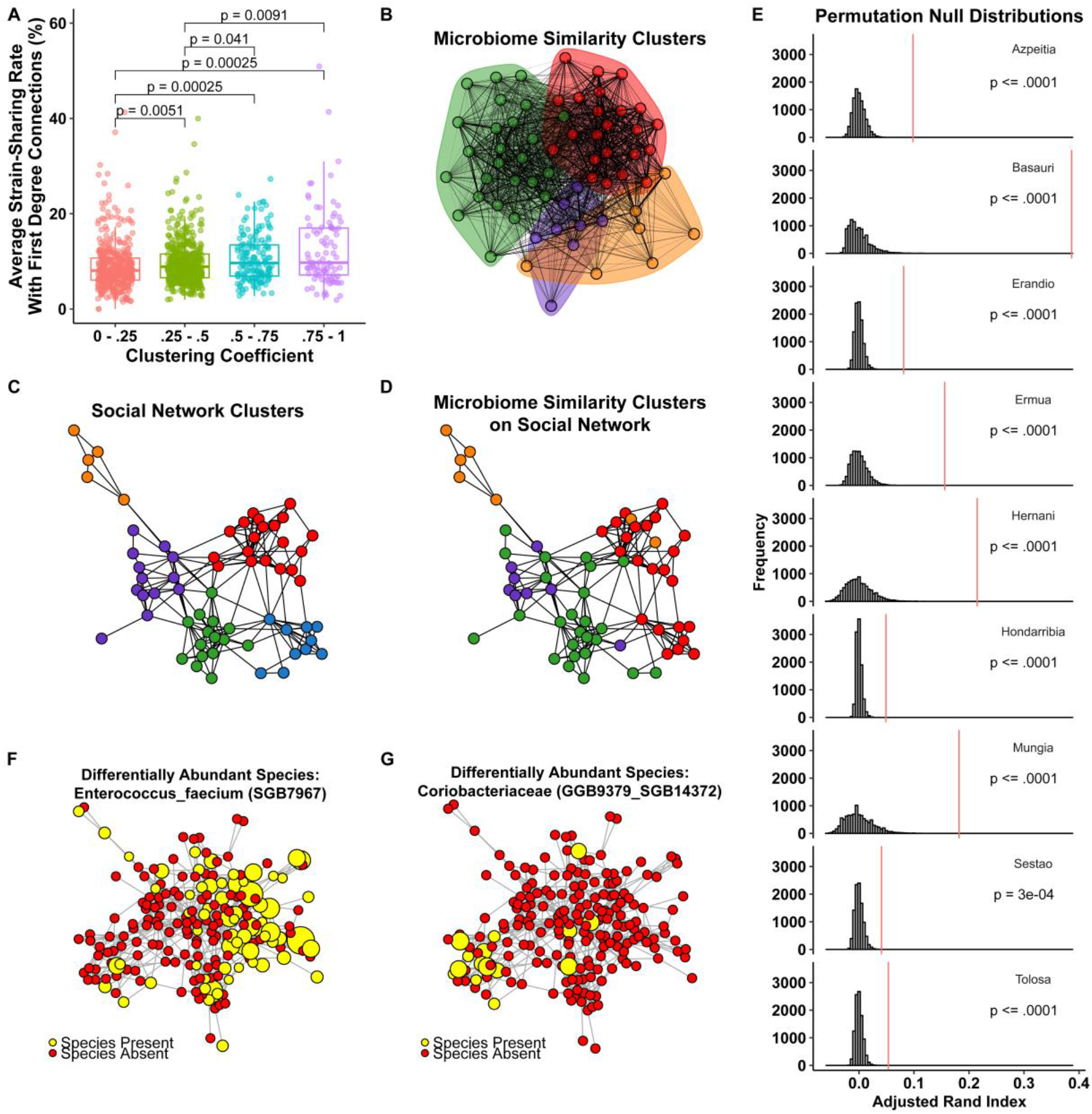
Social and Microbiome Strain Niches. **(A)** Individuals with a higher clustering coefficient are on average more similar to their first-degree connections. (**B)** Microbiome strain-sharing-rate Louvain clusters for the village of Basauri. Ties are weighted and sized according to the strain-sharing rate between the pair. **(C)** Social network Louvain clusters for the village of Basauri. **(D)** Microbiome cluster membership painted onto the social network. There is visual overlap between communities detected solely on the basis of shared microbe sets and communities detected based solely on social connections. **(E)** P-value distributions for clustering results. Histograms represent the null distribution of adjusted Rand index values from 10,000 microbiome permutations. All villages are highly significant, with the observed overlap metric represented by the vertical red line. **(F** and **G)** Examples of differentially abundant species within the village of Hondarribia. Nodes are scaled according to the log relative abundance of the species, with yellow indicating presence of the species and red indicating absence.

To observe this phenomenon at the whole-village-level scale, we identify both social and microbiome clusters using Louvain clustering,^36–38^ a method that can discern clusters in social network data^38–40^ and biological data^41,42^. If strain-sharing rates are significantly elevated within social network clusters, we would expect a large correspondence between social network clusters and clusters of microbially similar individuals. We form microbiome clusters based on the strain-sharing network within a village, with ties between people discerned simply by virtue of the extent to which they share microbiome strains, with edges weighted by the strain-sharing rate between constituent nodes (**Fig. 5B)**. Similarly, we form social clusters based on familial and social connections without weighting (**Fig. 5C**). On average, this method yields social clusters of 13 people with an average of 27 intra-cluster relationships; and it yields microbiome clusters of 24.5 people with an average intra-cluster strain-sharing rate of 7.5%. We can then paint the microbiome cluster membership onto the social network clustering and visualize the correspondence between social communities and microbiome strain communities **(Fig. 5D)**.

Across the villages, social clusters visually overlap with microbiome clusters (shown for just one village in **Fig. 5C** and **5D**). To statistically test this effect, we can evaluate the correspondence between social and microbiome cluster membership with the adjusted Rand index^43^, which is a measure of similarity between two data clusterings, with an adjustment for the chance grouping of elements. To observe the distribution of this statistic if there was independence between a host’s microbiome and their social network, we can compare our observed index to a microbiome permutation null, where we randomly swap every individuals microbiome in the village. We observe that social cliques correspond to microbial cliques at a highly significant rate in all nine villages (max *P* < .05) (**Fig. 5E**). Across 10,000 microbiome permutations, in only 1 village does a random permutation lead to higher overlap between social and microbiome clusters than the observed overlap (Sestao in **Fig. 5E**, where 3 out of 10,000 permutations have a higher overlap between clusterings).

The co-occurrence of strain-sharing clusters and social clusters suggests that certain species, in addition to strains, may also be differentially abundant in different social network niches. If social clustering reinforces within-group microbial sharing, we would expect different social clusters to have differentially abundant bacteria. To test this effect, we compared whether the relative abundance of each species differed across Louvain social clusters in each village with the Kruskal-Wallis test. After Benjamini-Hochberg multiple testing correction, we found 125 examples of species that were differentially abundant in different network communities out of 9,366 tests (Fig. S12 shows the p-value distributions of this test). **Fig. 5F and G** show examples of two species differentially abundant in different network regions of an illustrative village; *Enterococcus faecium (SGB7967)* is highly abundant in the right-hand side of the network, whereas *Coriobacteriaceae (SGB14372)* is highly abundant in the bottom left region of the village.

## Discussion

Using detailed social network mapping and strain-level microbiome genomics in isolated Honduras villages, we find a substantial correspondence between social structure and microbiome sharing beyond familial or even household relationships. The amount of strain-sharing appears to be modulated by the nature of the social relationships. More intimate relationships share more strains, and strain-sharing rates monotonically increase based on the frequency a pair shares meals or free time together. The strain-sharing rate was the strongest predictor of social relationships, which suggests that when entering a village and knowing nothing about the social arrangements, there is more predictive power in knowing the microbiome distribution than knowing a sociodemographic feature such wealth, religion, or education. We also observe significantly elevated strain-sharing levels out to a social horizon of third-degree connections. And host network position, whether central or peripheral, moderates egocentric exposure to the microbial metacommunity within the villages; more socially isolated people tend to be more microbially isolated. Overall, the intricate groundwork provided by the social network structure of human populations provides a set of niches within which microbes can thrive or spread.

We are unable to distinguish direct transfer of strains from indirect transfer (e.g., via unobserved social interactions that we were unable to measure), nor can we infer the directionality of any transfer between two people seen to share a strain. We are also unable to fully distinguish shared environment from transmission, though the genetic specificity of strains and the distinction between villages or far-flung households does suggest actual transmission, especially in light of the human-host specificity of some transmitted species^44–47^. Strain-level resolution helps shed light, to some extent, on the idea that similar microbial species seen within members of the same household may be based not only on a modulation by similar environmental conditions or shared genetics, but also on spread between individuals. Our ability to also find strain-sharing among people who are not genetically related and do not reside in the same household but who are known to interact bolsters this conclusion.

A prior study of 287 people in five villages in Fiji was able to document strain sharing between spouses, household members, and a subset of other social interactions (up to five other people with whom they “spent the most amount of time” (a total of 489 unique social interactions outside households were identified)^11^. Another recent study examining 7,646 individuals from 31 villages in 20 countries observed that, on average, the strain-sharing rate for the gut microbiome for non-cohabiting adults within the same village generally was 8%^24^. Our estimate of this parameter was 3.9%. But, since we actually mapped so many types of social relationships (and not just familial or same-household ties) among essentially all the inhabitants of each village, we can be more sure than prior studies about whether village co-residents do or do not, in fact, interact with each other. That is, our estimate of pairs of people who are simply village co-residents includes only people who do *not,* in fact, interact; this likely explains the difference in findings. Social network mapping at the level of detail, comprehensiveness, and scale that is present here has not previously been available for microbiome analyses.

Findings with oral or skin microbiomes might be different. Oral strain-sharing rates are known to be higher than seen with the gut; for instance, among cohabiting individuals, prior work has shown that the median strain-sharing rates are 12% and 32% for gut and oral microbiome respectively^24^. And oral microbiome strains are known to be more easily transmitted, via saliva^11,33^. Future work could involve such specimens. Future research could also include longitudinal information, with suitable temporal spacing, ascertaining both the social network interactions and the microbiome status of all villagers, evaluating any change in microbes and the sharing of species or strains that might accompany any change in social network niches.

The Baas-Becking hypothesis suggests that “everything is everywhere, but the environment selects,” and it is provocative to consider whether there are particular social niches within the social network fabric of human connections that are more conducive to certain species rather than to others^48^. Our social networks may provide the extended niches in which microbes can thrive. And being exposed to the microbes of others within one’s social group can have either benefits or drawbacks. For example, if a person becomes ill or takes antibiotics and loses some of the helpful microbes in their body, they could perhaps regain them through exposure to the beneficial microbes of others. Conversely, people with a diversity of social connections may also be more likely to be exposed to harmful microbes circulating within a network.

Finally, using both observational and experimental methods, diverse phenomena have been shown to spread interpersonally, including phenotypes such as obesity and depression^49–51^. To the extent that the microbiome can be associated with physical or mental states (and there is increasing evidence for this^52^), then any spread of the microbiome via biological contagion may partly explain the ostensible spread of other attributes via social contagion. Groups of interconnected people may share phenotypes not only because of shared genes or transmitted behaviors, but also because of shared microbes.

## METHODS

### Network Construction

Village-level networks were mapped with standard “name generators” for the whole village. After a photographic census (of all adult residents) was taken for each village, we conducted the main network survey in each village, including a detailed, hour-long survey^17^, incorporating demographic and health measures, as well as a battery of name generators with which respondents identified relevant social relationships (friends, family members, etc.) through names and images shown in our TRELLIS software (available at trellis.yale.edu)^53^. All the name generator questions are listed in Table S1. Name generators used to ascertain social ties included standard questions such as “with whom do you spend free time?” and “with whom do you discuss personal or private matters?” as well as questions about closest friends.

For questions in which a pair may report different levels of the same variable, such as greeting type or the amount of free time, we symmetrize the variables as follows: For greeting type, we report the greeting type involving the most physical contact. For the frequency of free-time and shared meals between a pair, we symmetrize by choosing the response between the pair that indicates more frequent contact. We symmetrized all other responses at the relationship level (i.e., when two individuals nominate each other as a close friend, the relationship is only counted once). When calculating degree distributions, centralities, and clusterings, we simplified our networks to remove multiplexity (i.e., we concatenated all ties between a pair).

Social network graphs were analyzed and geodesic distances and centrality measures were calculated with igraph (v1.3.5)^54^ and plotted with the Fruchterman-Reingold algorithm. We also collected GPS coordinates for all households. To protect the anonymity of our study region, villages were renamed to random town names from another country.

### Gut Microbiome Sample Collection, Library Preparation, and Sequencing

Participants were instructed on how to self-collect the fecal samples using a training module and asked to return samples promptly to the local team. Samples were then stored in liquid nitrogen at the collection site and moved to a −80 C° in Copan Ruinas, Honduras. Samples were shipped on dry ice to the United Stated of America and stored in −80 C° freezers.

Stool material was homogenized using TissueLyzer from Quigen and the lysate was prepared for extraction with the Chemagic Stool gDNA extraction kit (Perkin Elmer) and extracted on the Chemagic 360 Instrument (Perkin Elmer) following the manufacturer’s protocol. Sequencing libraries were prepared using the KAPA Hyper Library Preparation kit (KAPA Biosystems). Shotgun metagenomic sequencing was carried out on Illumina NovaSeq 6000. Samples not reaching the desired sequencing depth of 50Gbp were resequenced on a separate run. Raw metagenomic reads were deduplicated using prinseq lite (version 0.20.2^55^) with default parameters. The resulting reads were screened for human contamination (hg19) with BMTagger and then quality filtered with Trimmomatic^56^ (version 0.36, parameters ILLUMINACLIP: nextera_truseq_adapters.fasta:2:30:10:8:true SLIDINGWINDOW: 4:15 LEADING: 3 TRAILING: 3 MINLEN: 50). This resulted in a total of 1,188 samples (with an average size of 8.6×10^7^ reads).

### Species-level and Strain-level Profiling

Species-level profiling was performed using MetaPhlAn 4 using the Jan21 database and default parameters. Strain-level profiling was performed for a subset of species present in at least 50 samples using StrainPhlAn 4^57^ with parameters ‘--marker_in_n_samples 1 -- sample_with_n_markers 10 --phylophlan_mode accurate’. This resulted in a total of 682 SGBs and 183,195 profiled strains. The StrainPhlAn ‘strain_transmission.py’ script was used to assess transmission events using the produced trees which yielded a total of 336,710 identified events. For a robust calculation, strain-sharing rates were calculated only for pairs sharing at least 10 SGBs.

### Statistical Analyses

All statistical analyses were performed in R (v.4.1.3) with additional packages ggpubr (v0.5.0) and ggplot2 (v3.4.0). Correction for multiple testing (Benjamini–Hochberg procedure, marked *P*_adj_) was applied when appropriate, and significance was defined at *P*_adj_ < 0.05. All tests were two-sided except where otherwise specified. All egocentric regressions involved linear mixed effects models with this general specification:

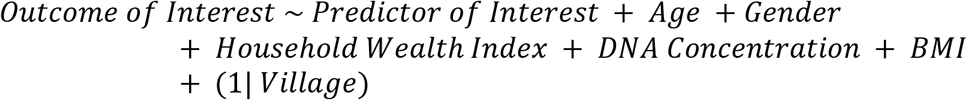

Mixed effects models were created with the lmertest package (v3.1.3)^58^.

Figure 1 was generated using Python (v3.9.7). The scikit-bio package (v0.5.6) was used for calculating the centered log-ratio transformation of the microbiome species relative abundances after all zero values were replaced with the small value of 5×10^-6^, which was half the value of the original smallest non-zero value before replacement.

The sklearn package (v1.0.1) was then used to compute the t-distributed stochastic neighbor embedding (t-SNE) lower-dimensional representation of the centered log-ratio transformed data with a perplexity parameter of 5 and random seed 0. The 2D t-SNE representation was then visualized using the matplotlib package (v3.4.2), coloring points by village membership.

### Network Predictions

Mixed effect logistic regressions were used for out-of-sample network predictions. Class-balanced data sets were constructed by down-sampling the number of unrelated pairs to equal the number of related pairs. We analyzed our model with threefold cross-validation. Predictions from the three separate test sets were combined. Receiver operating characteristic (ROC) curves were constructed from the average of five sets of threefold cross-validation. ROC Curves and confidence intervals were calculated with the pROC package (v1.18.0)^59^ and logistic regression models were constructed with the lmertest package (v3.1.3) with the binomial family link function and a random slope per village. The predictive model including all covariates was specified by:

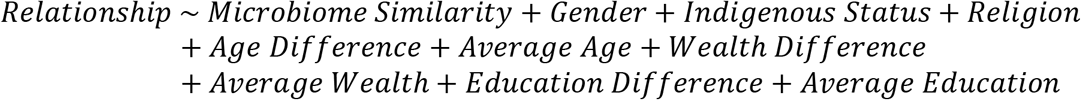

Variable importance metrics were calculated based on the permutation feature importance metric. The permutation feature importance is defined to be the decrease in a model score when a single feature value is randomly shuffled^34^. This procedure breaks the relationship between the feature and the target; thus, the drop in the model score is indicative of how much the model depends on the feature. Variable importance metrics were analyzed after 1000 random permutation of each feature. VIF values were calculated to ensure the reliability of results against colinearity of variables and were all low (<2).

### Microbiome Null Permutations

Microbiome null permutations create a null distribution of strain-sharing rates between any two people while accounting for network structure. Under the null hypothesis that a host’s microbiome composition and social network are independent, we can sever their relationship by randomly permuting the microbiome of every host in the village and recalculating metrics of interest, e.g., strain-sharing by degree or clustering Rand indices. This ensures that the inherent structural pattern of the network remains the same, but the node values are randomized. This allows us to observe the distribution of our statistics if the human microbiome is fostered independently of any host social interactions.

Village-wide microbiome permutations were used to calculate null distributions for the strain-sharing rate by geodesic distance and for the clustering results. For relationship-specific permutations in Fig. S1, permutations at the relationship level were taken instead of full village permutations. The observed distribution of relationship-specific sharing was compared to the distribution of sharing observed when that specific relationship tie was permuted, for example comparing the sharing between someone and their friend versus someone and 100 random individuals’ friends in the same village. For the inherently gendered relationships of spouse and mother/father to the child, we accounted for the gender of the ego; but for all other relationships which are not inherently gendered (e.g., free time), we did not.

### Microbiome and Social Clustering

We use the Louvain method as implemented in the igraph package to cluster our participants along social and microbiome lines. Louvain clustering is based on greedy modularity optimization. Modularity is a scale value between −0.5 (non-modular clustering) and 1 (fully modular clustering) that measures the relative density of edges inside communities compared to edges outside communities. Optimizing this value theoretically results in the best possible grouping of the nodes of a given network. In cases where a pair shared too few SGBs to calculate a robust strain-sharing rate (<10), a strain-sharing rate of 0% was imputed to allow for proper weight-based clustering. The adjusted Rand index was calculated with the mclust package (v6.0.0)^60^.

For testing species differential abundance across network communities with the Kruskal-Wallis test, robustness checks ensuring that each social cluster has more than 5 people and the species is present in more than 5 people in the village were performed, and cases where this criterion was not met were excluded.

## Supporting information

Supplementary Informations

## Acknowledgments

We thank all the study participants in Honduras. We thank Rigoberto Matute Juarez, Jose Eduardo Gámez, and Eduardo Jose Urrea Carbajal for coordinating the field work; Rennie Negron, Liza Nicoll, and Thomas Keegan for their support on field operations, data collection, and administrative support; and YCGA (Yale Center for Genomic Analysis) for sequencing the metagenomic libraries; and Qiaojuan Shi for processing the specimens and handling the extractions. We thank Michael Baym and Mark Gerstein for helpful comments.

This work was supported by the NOMIS Foundation, with additional support from Schmidt Futures, the Pershing Square Foundation, and the Rothberg Catalyzer Fund.

## Author contributions

Conceptualization: JP, FB, SS, MA, DP, IB, and NAC; Methodology: JP, FB, SS, MA, DP, IB, and NAC; Data Collection: FB, MA, IB, and NAC; Statistical Analysis: JP, FB, SS, MA; Funding acquisition: NAC; Supervision: IB, NAC; Writing: JP, FB, SS, MA, DP, IB, and NAC.

## Data and materials availability

The codebase for this work will be posted at a public URL.

