## Supplementary Informations for "Detailed Social Network Interactions and Gut Microbiome Strain-Sharing Within Isolated Honduras Villages"

**Supplementary Information**

**Figure List:**


1.) **Strain-Sharing by Relationship Permutation P-values**
2.) **Species-level Sharing (Bray-Curtis)**
3.) **Species-level Sharing (Jaccard)**
4.) **Non-Kin Different-House Strain-Sharing**
5.) **Strain-Sharing by Physical versus Non-Physical Greetings**
6.) **Parent-to-Child Microbiome Sharing**
7.) **Microbiome Sharing by Gender**
8.) **Species Relationship Prediction Model**
9.) **Strain-Sharing Relationship Prediction Model Permutation Feature Importance**
10.) **Species-Level Sharing by Geodesic Distance**
11.) **Decline in Average Microbiome-Sharing with Network Size**
12.) **Species Niches P-Value Distributions**


**Table List:**


1.) **Selected Name Generator Questions**

**Figure 1: Strain-Sharing by Relationship Permutation P-values**

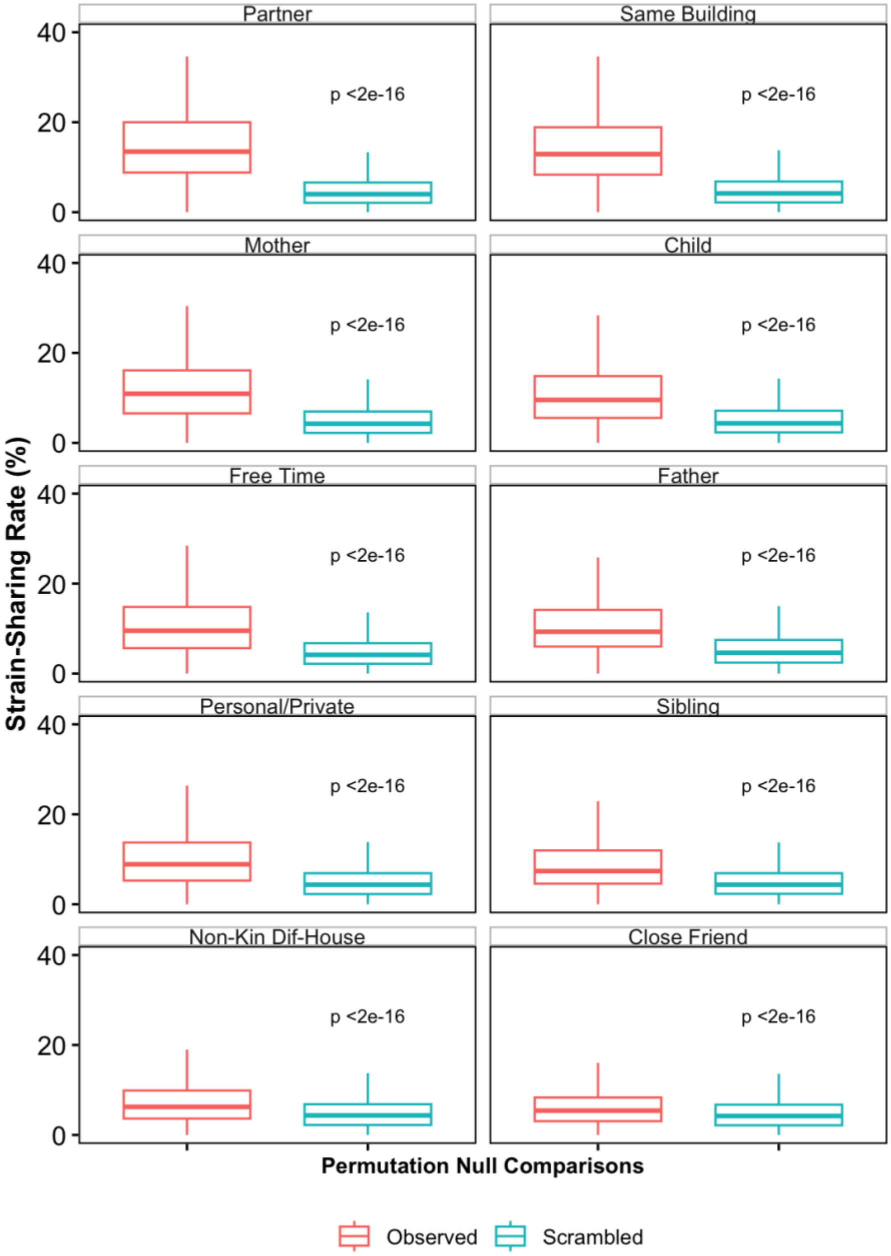


Observed strain-sharing rates for each relationship compared to 100 draws from a within-village relationship permutation. For example, the red distribution would contain the strain-sharing rate between an individual and their mother, and the blue distribution contains the strain-sharing rate for the same individual with 100 random mothers in the same village. All observed relationships have a significantly higher strain-sharing rate than the scrambled networks, with the adjusted P-value reported in each figure (two-sided Wilcoxon rank-sum tests).

**Figure 2: Species-level Sharing (Bray-Curtis)**

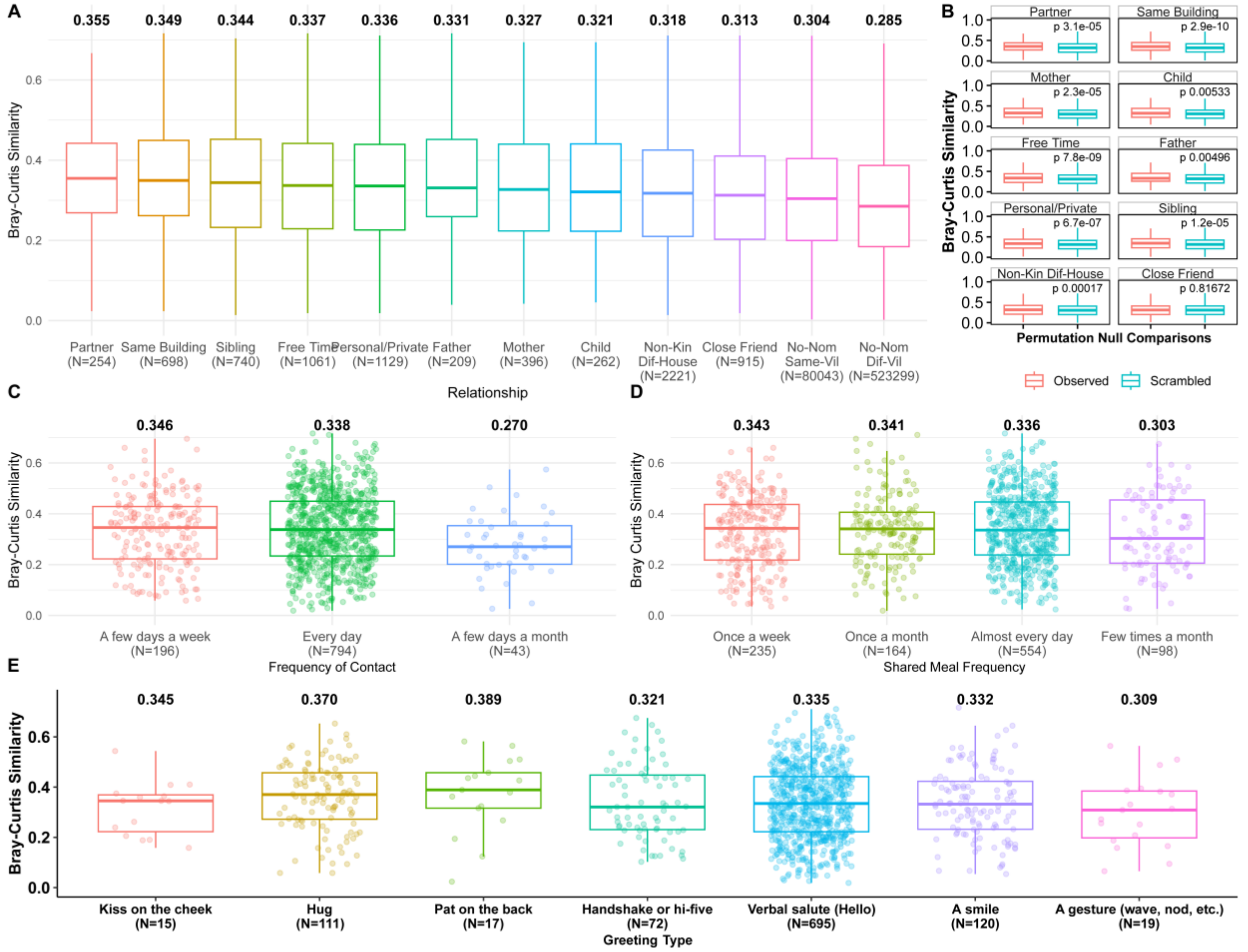


**(A)** The distribution of Bray-Curtis similarity based on relationship type. The final two boxes contain the strain-sharing rates between individuals living in the same village without a nominated relationship, and all pairs of individuals living in different villages, respectively. Median values for each distribution are printed at the top of each box. **(B)** Observed Bray-Curtis similarity for each relationship compared to 100 draws from a within-village relationship permutation. All observed relationships, except for close friends, have a significantly higher Bray-Curtis similarity than the scrambled networks, with the adjusted P-value reported in each figure (two-sided Wilcoxon rank-sum tests). **(C)** Bray-Curtis similarity by how often a pair spends free time together. **(D)** Bray-Curtis similarity by how often a pair shares meals together. **(E)** Bray-Curtis similarity by greeting type.

### Figure 3: Species-level Sharing (Jaccard)

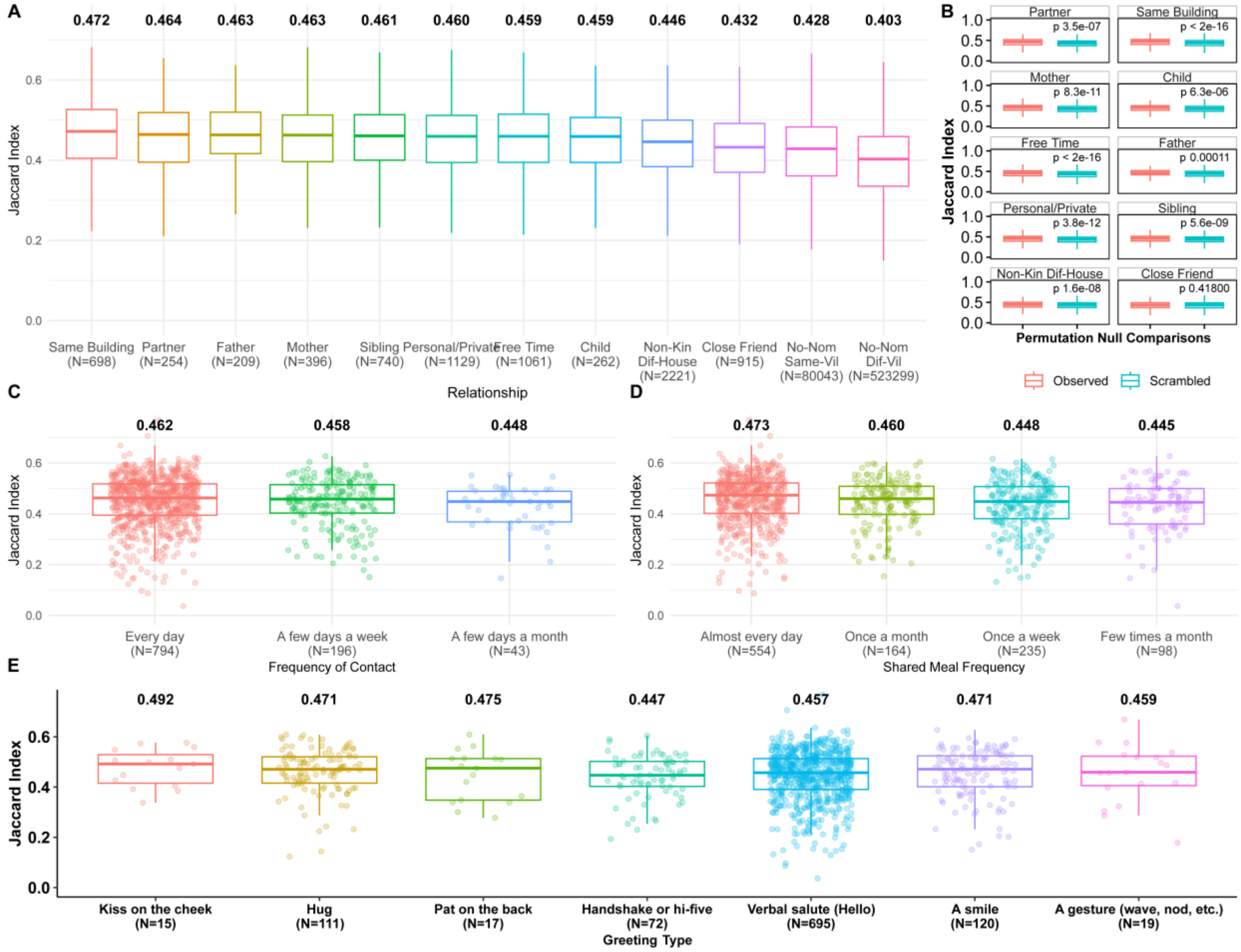


**(A**) The distribution of Jaccard similarity based on relationship type. The final two boxes contain the strain-sharing rates between individuals living in the same village without a nominated relationship, and all pairs of individuals living in different villages, respectively. Median values for each distribution are printed at the top of each box. **(B)** Observed Jaccard similarity for each relationship compared to 100 draws from a within-village relationship permutation. All observed relationships, except for close friends, have a significantly higher Jaccard similarity than the scrambled networks with the adjusted P-value reported in each figure (two-sided Wilcoxon rank-sum tests). **(C)** Jaccard similarity by how often a pair spends free time together. **(D)** Jaccard similarity by how often a pair shares meals together. **(E)** Jaccard similarity by greeting type.

**Figure 4: Non-Kin Different-House Strain-Sharing**

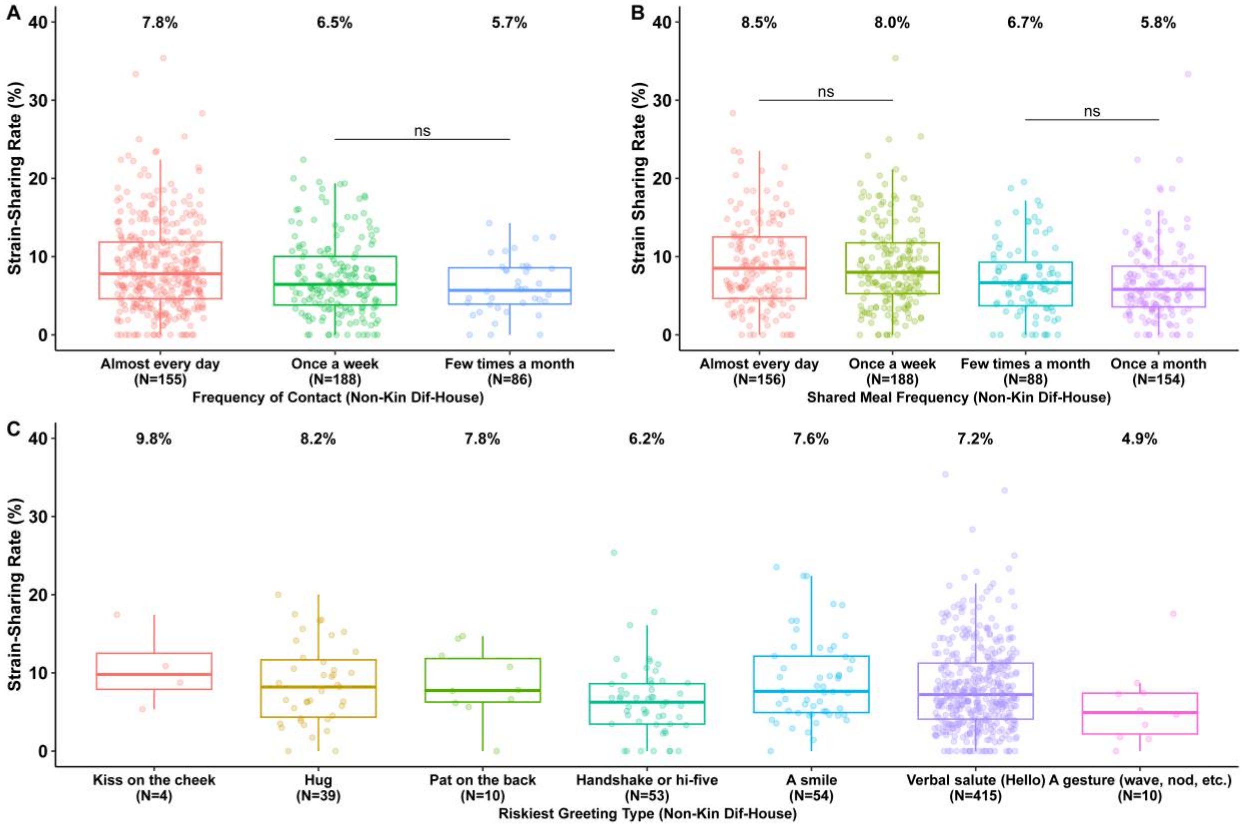


**(A)** Strain-sharing amongst non-kin different-house relationships by frequency of free-time contact. **(B)** Strain-sharing amongst non-kin different-house relationships by frequency of shared meals. **(C)** Strain-sharing amongst non-kin different-house relationships by greeting type.

**Figure 5: Strain-Sharing by Physical vs Non-Physical Greetings**

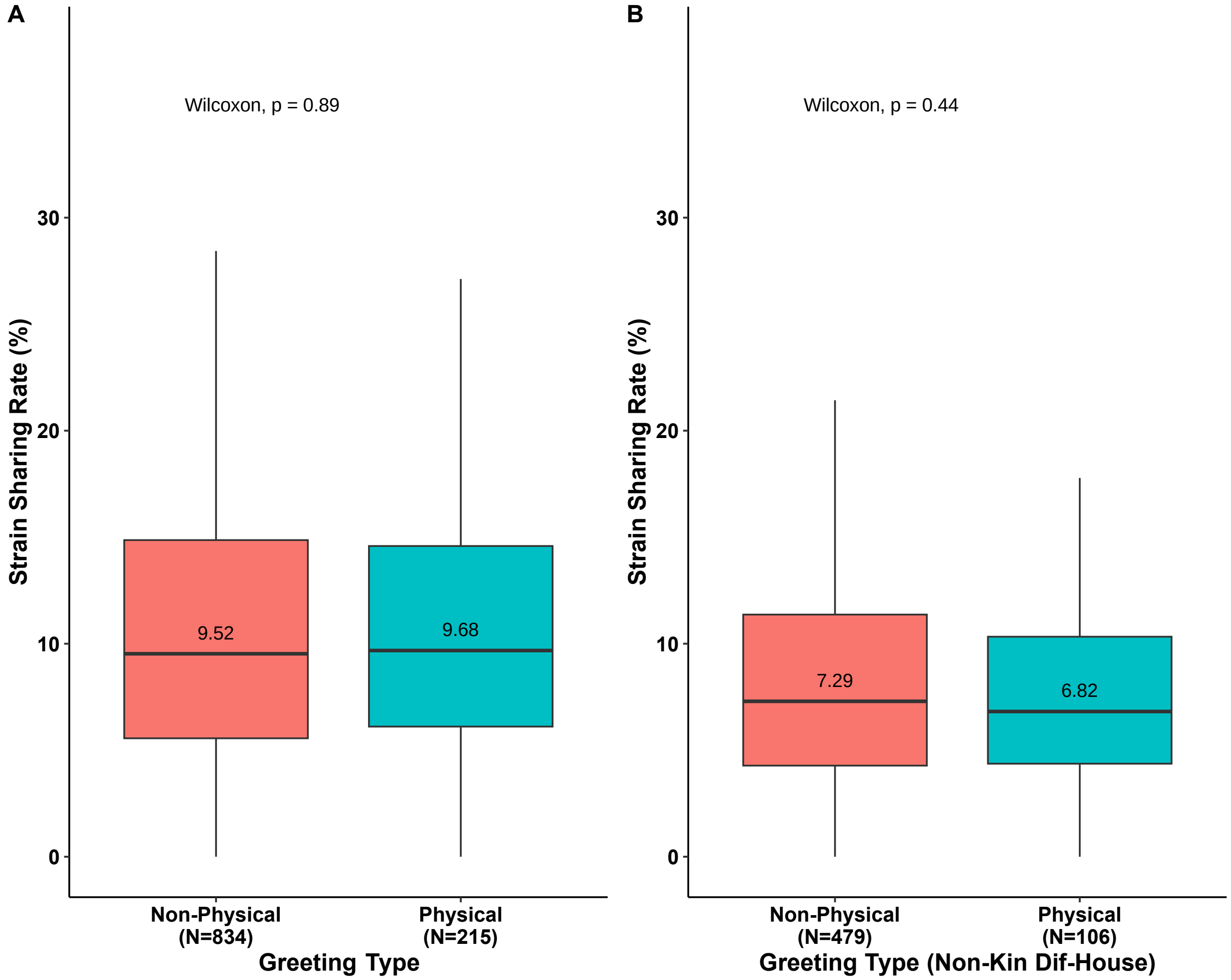


**(A)** Strain-sharing rates by dichotomized greeting type (physical versus non-physical) for all relationships. **(B)** Strain-sharing rates by dichotomized greeting type (physical versus non-physical) for all non-kin different-house relationships.

**Figure 6: Parent-to-Child Microbiome Sharing**

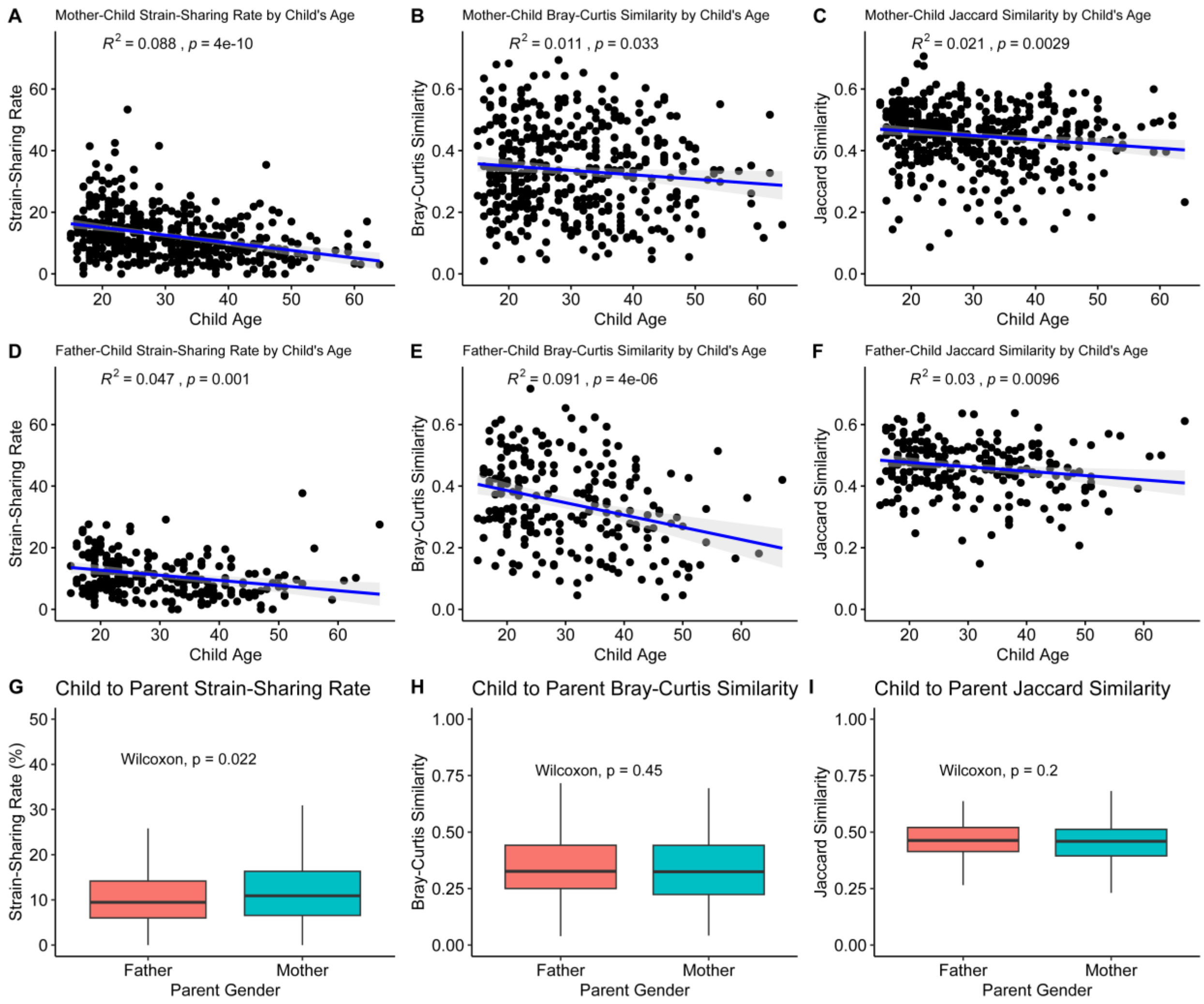


Parent-to-child microbiome strain-sharing rates decrease as children age and gain exposure to different environments. The trend is seen for both mothers (top row) and fathers (bottom row) based on strain-sharing rates **(A** and **D)**; species Bray-Curtis similarity **(B** and **E)**; and species Jaccard index **(C** and **F)**. On average, mothers have a higher strain-sharing rate to their children than fathers **(G)** but this trend is not observed at the species level **(H** and **I**).

### Figure 7: Microbiome Sharing by Gender and Relationship

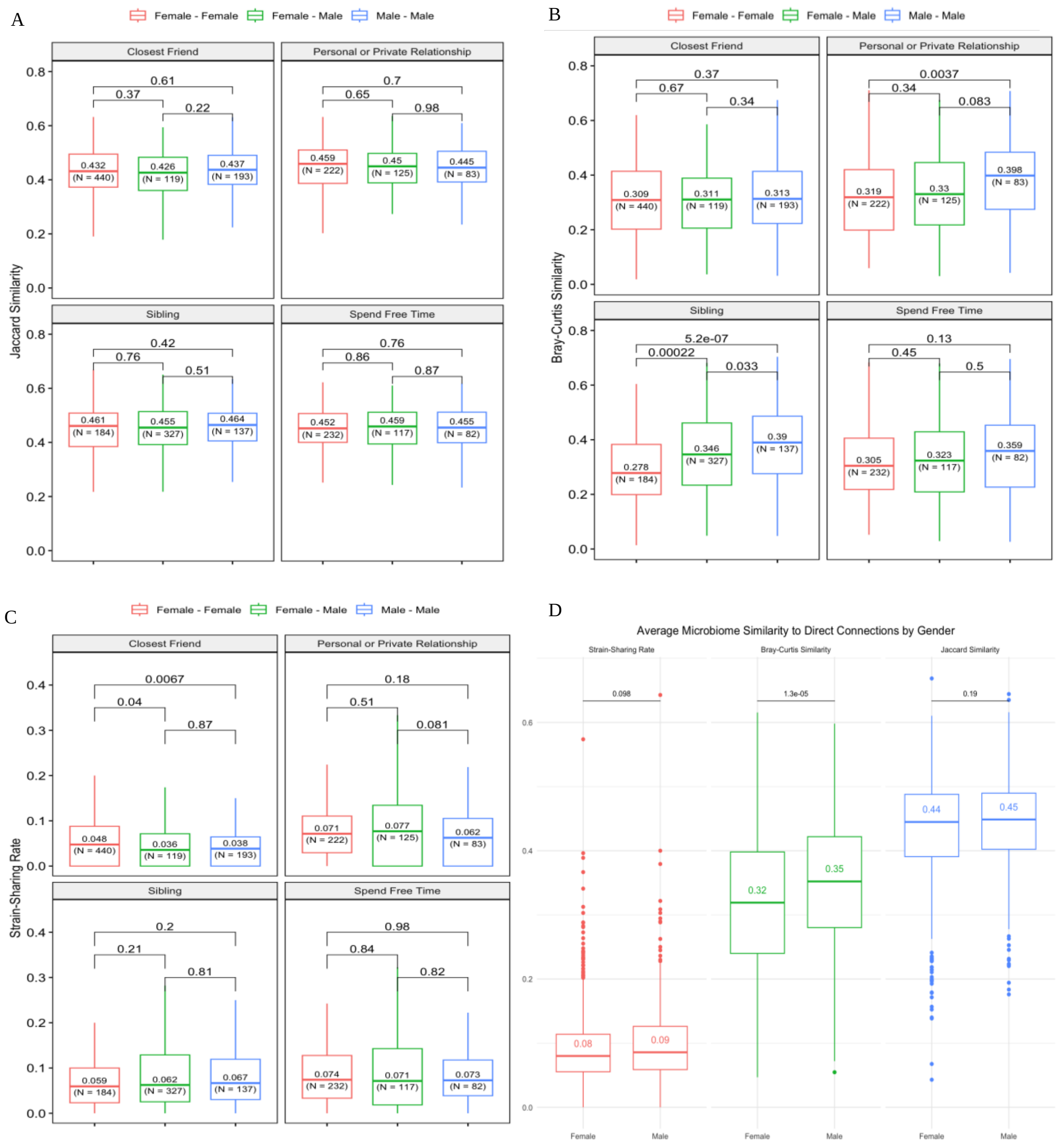


Microbiome sharing stratified by relationship and genders involved based on **(A)** Jaccard similarity, **(B)** Bray-Curtis similarity, and **(C)** Strain-sharing rate. **(D)** Microbiome similarity to direct connections based on host gender. Men tend to have a higher Bray-Curtis similarity to their direct connections than women. P-values based on two-sided Wilcox Rank Sum tests are shown at the top of each comparison.

**Figure 8: Species Relationship Prediction Model**

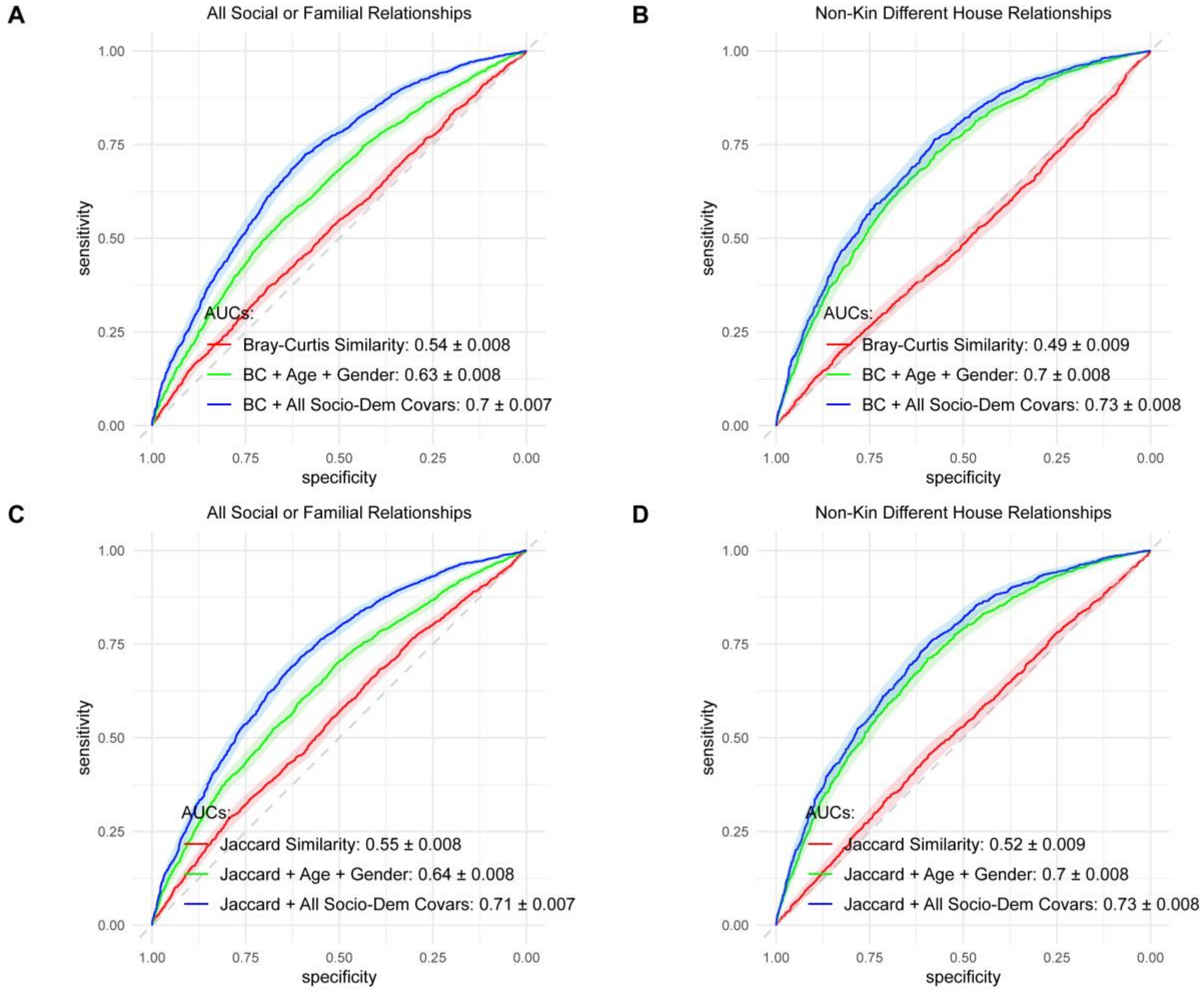


Social network prediction model results using species level data with Bray-Curtis (top row) and Jaccard similarity (bottom row) for all relationship types (left column) and non-kin different-house relationships (right column).

**Figure 9: Strain-Sharing Relationship Prediction Model Permutation Feature Importance**

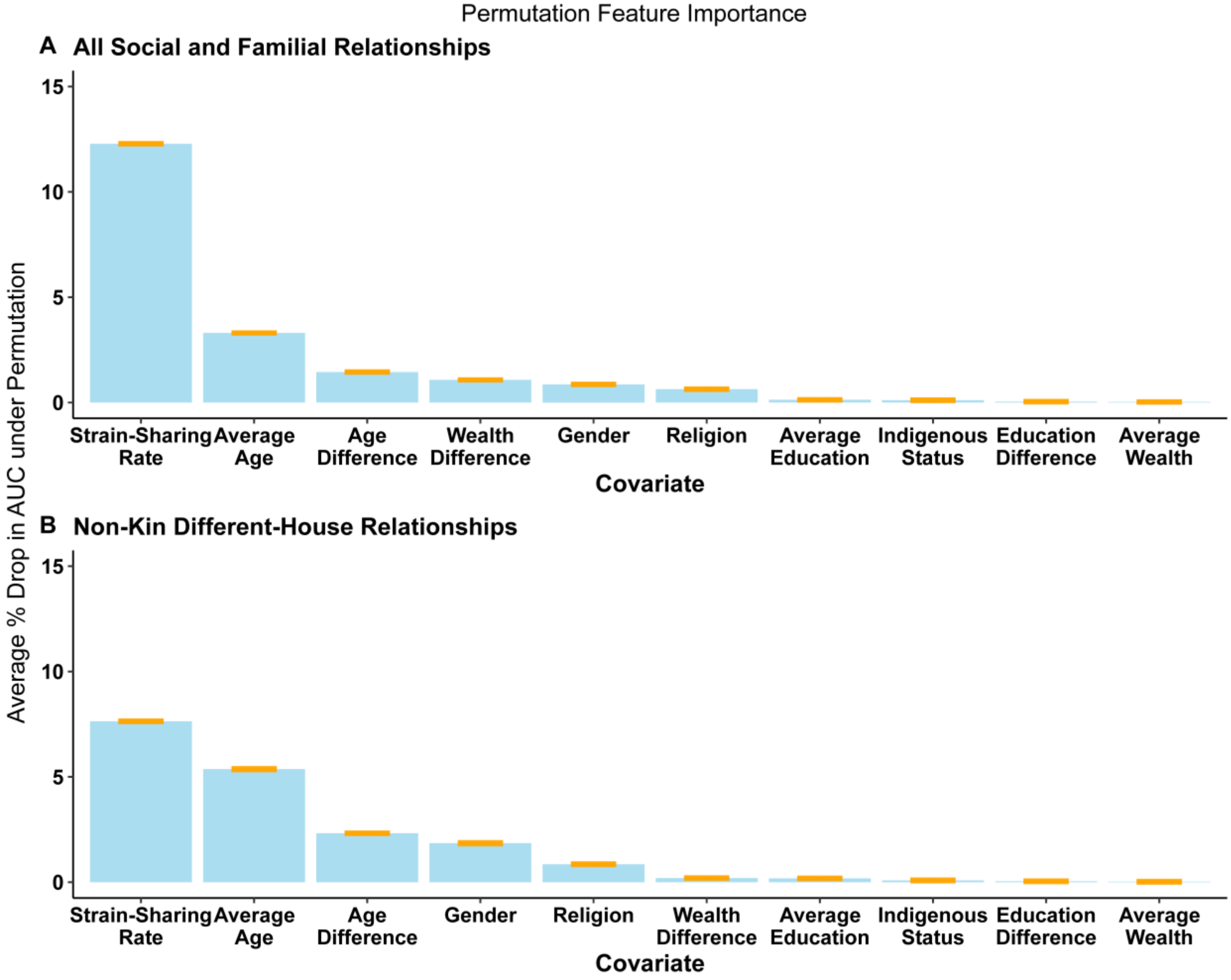


Permutation feature importance results for all relationships **(A)** and non-kin different-house relationships **(B)**. In both models, the strain-sharing rate is the strongest predictor of a relationship. Orange bars at the top of each plot indicate 95% confidence intervals for the drop in model score.

### Figure 10: Species-Level Sharing by Geodesic Distance

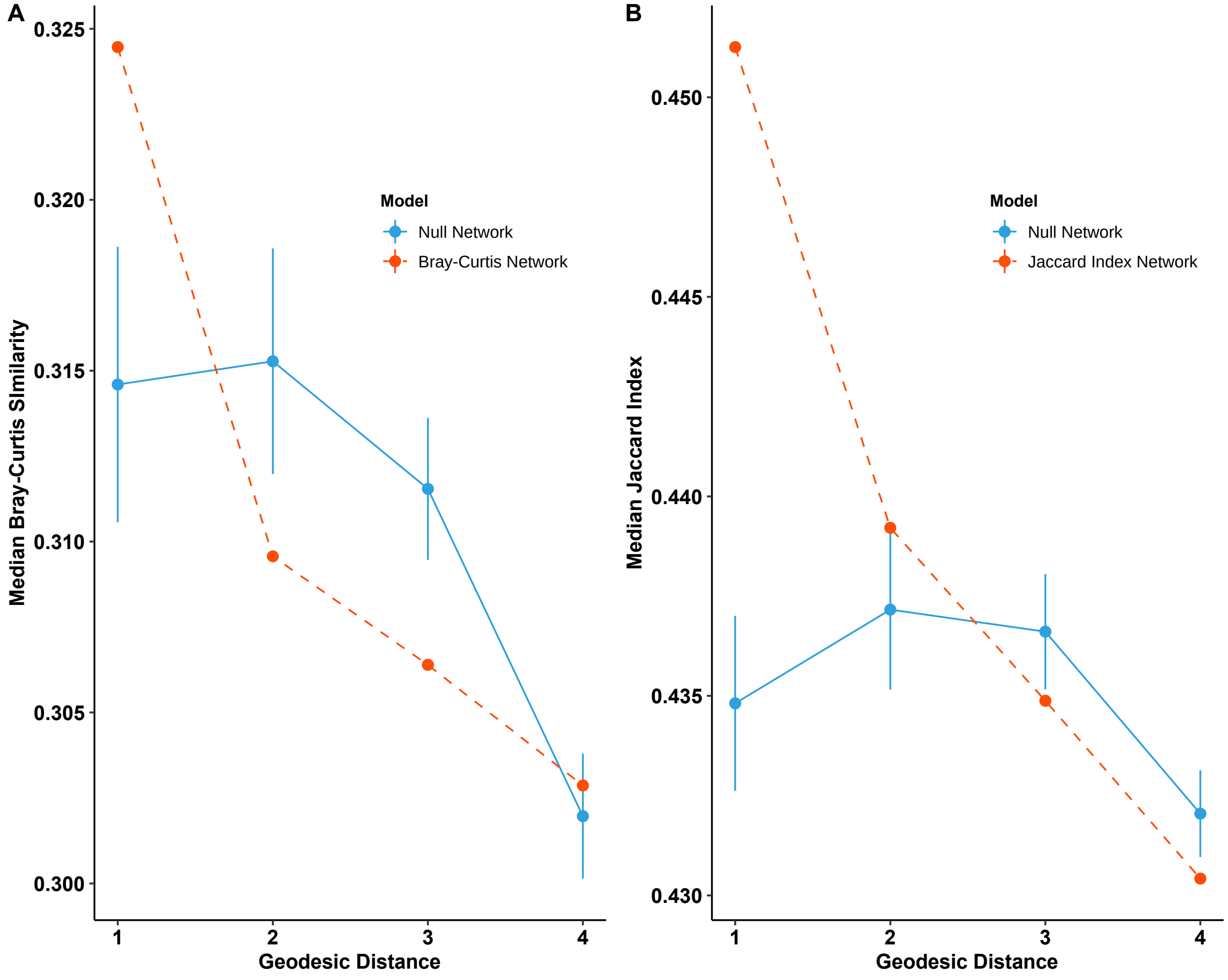


Average Bray-Curtis similarity **(A)** and Jaccard similarity **(B)** by geodesic distance in the social network. Bray-Curtis similarity is only significantly elevated for first-degree connections, while Jaccard similarity is elevated for first and second degree connections.

**Figure 11: Decline in Average Microbiome-Sharing with Network Size**

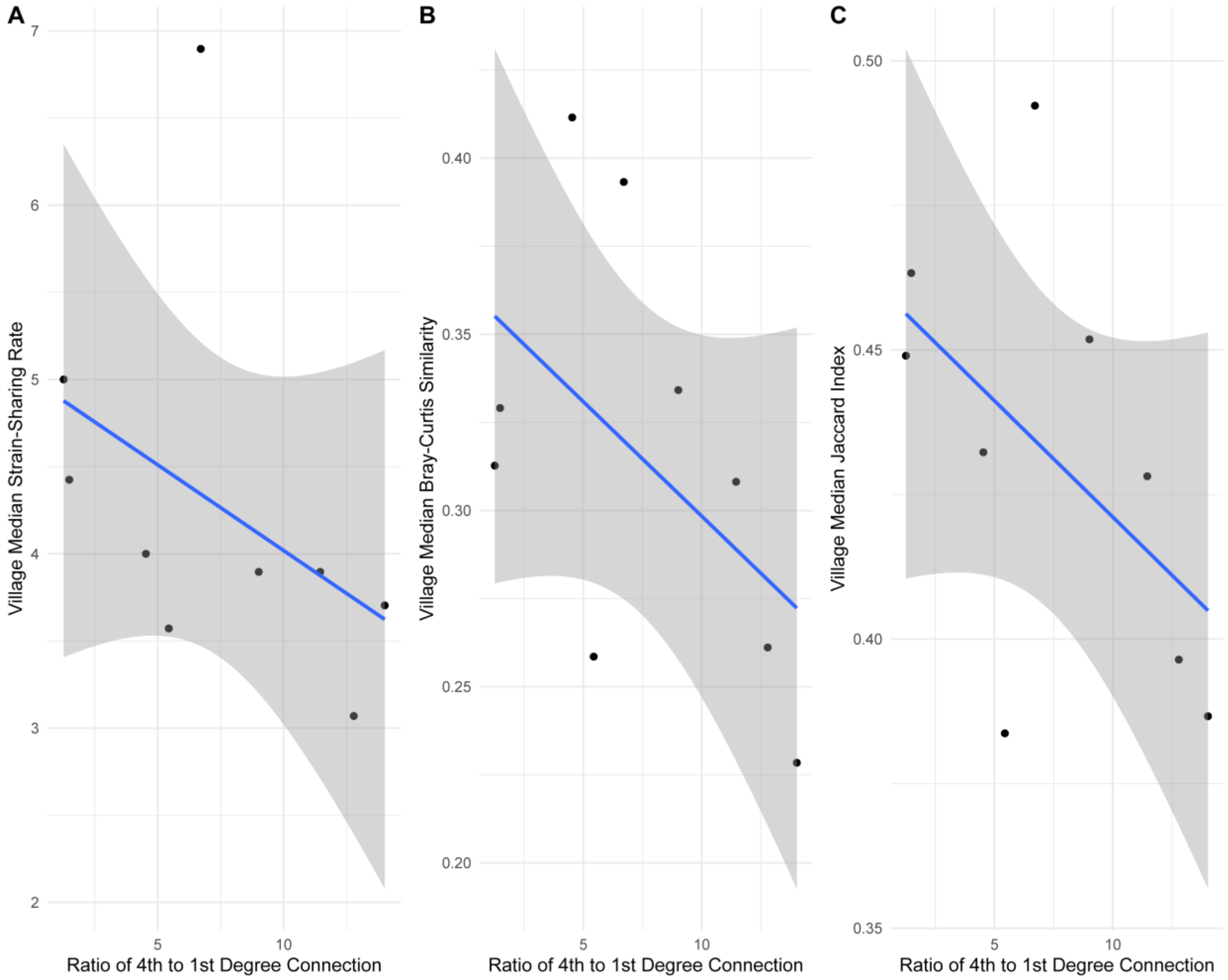


As the ratio of fourth to first degree connections increases, the average level of microbiome sharing amongst all individuals in a village tends to decrease. Consequently, the null distributions for microbiome-sharing by geodesic distance tend to dip downwards as larger villages have a higher ratio of fourth degree connections. Larger villages have higher village-level microbial diversity.

**Figure 12: Species Niches P-Value Distributions**

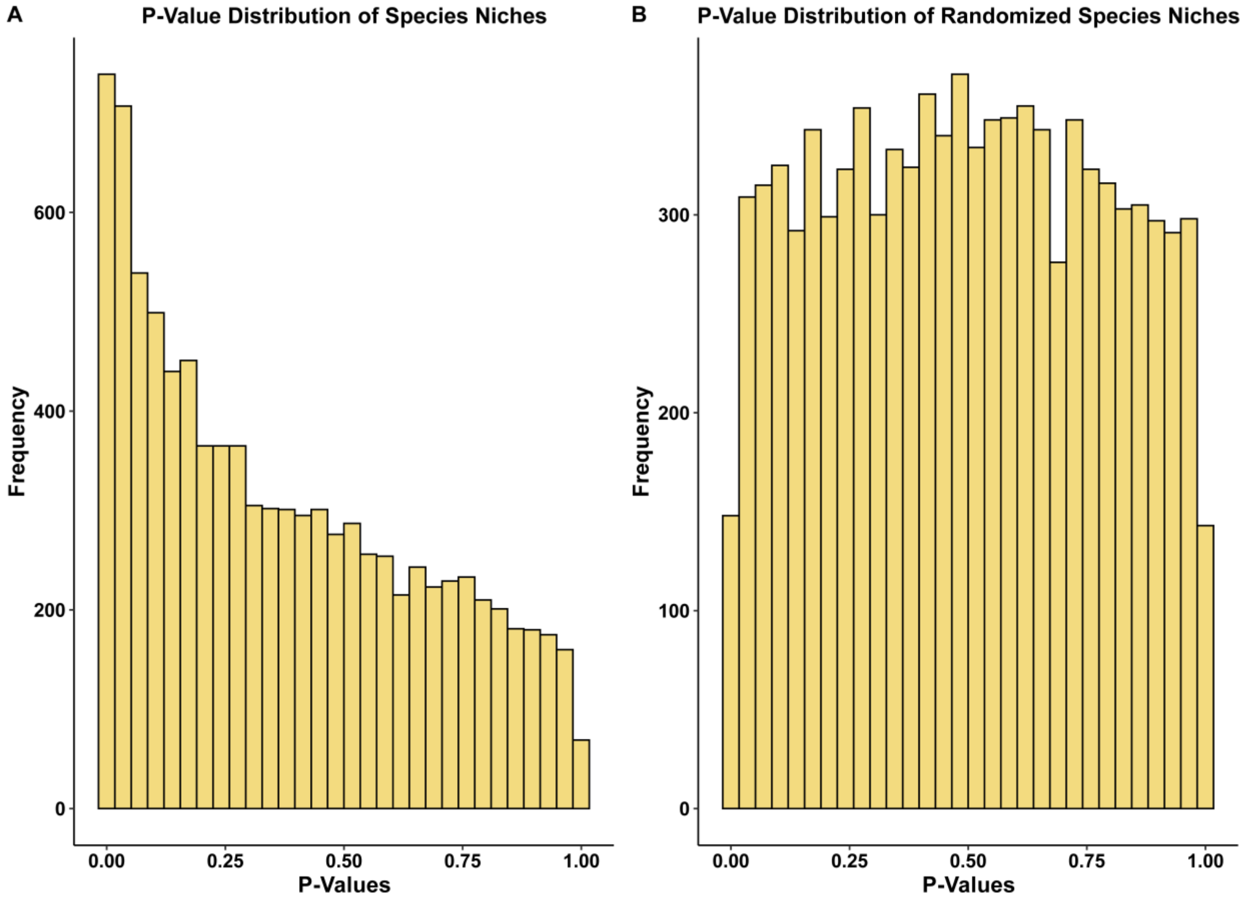


(**A**) Distribution of unadjusted p-values for the Kruskal-Wallis test for the differential abundance of species across network communities. The distribution is highly left skewed, indicating significant species clustering, whereas (**B)** under the null hypothesis that species are randomly distributed amongst village members, the distribution is uniform.

**Table 1: Name Generators to Map Networks**

| Kin Ties | General Ties (Kin and Non-Kin ) |
|---|---|
| - Does your mother live in this town? What is the name of your mother?<br>- Does your father live in this town? What is the name of your father?<br>- How many siblings do you have? How many are brothers? How many are sisters?<br>- How many of these siblings are over the age of 12 and currently living or working in this village? What are the names of your siblings over the age of 12 that live or work here?<br>- Do you have any children who don't live with you but do live in this village, over the age of 12? What are their names?<br>- Are you married or living in a civil union? What is the name of your partner? | - Who do you trust to talk to about something personal or private?<br>- With whom do you spend free time?<br>- Besides your partner, parents or siblings, who do you consider to be your closest friends? |
